# Pharmacological rescue of tumor intrinsic STING expression and immune response in LKB1-mutant lung cancer via the IAP-JAK regulatory axis

**DOI:** 10.1101/2021.09.17.460294

**Authors:** Changfa Shu, Rui Jin, Qiankun Niu, Danielle Cicka, Sean Doyle, Alafate Wahafu, Dacheng Fan, Xi Zheng, Yuhong Du, Andrey A. Ivanov, Deon B Doxie, Kavita M Dhodapkar, Jennifer Carlisle, Taofeek Owonikoko, Suresh Ramalingam, Gabriel Sica, Madhav V Dhodapkar, Wei Zhou, Xiulei Mo, Haian Fu

## Abstract

Harnessing the power of the immune system to treat cancer has become a core clinical approach. However, rewiring of intrinsic circuitry enables tumor cells to escape immune attacks, leading to therapeutic failure. Pharmacological strategies to reverse tumor genotype-dictated therapeutic resistance are urgently needed to advance precision immunotherapy. Here, we identify antagonists of Inhibitor of <u>A</u>poptosis <u>P</u>rotein (IAP) as potent sensitizers that restore immune-dependent killing of LKB1-mutant lung cancer cells. Mechanistic studies reveal an LKB1-IAP-JAK trimolecular complex that bridges the LKB1-mutant genotype with IAP-dependency and a STING-deficiency-mediated immune resistance phenotype. Ultimately, inhibition of IAP re-establishes JAK-regulated STING expression and DNA sensing pathway as well as enhanced cytotoxic immune cell infiltration and selective immune-dependent anti-tumor activity in an LKB1-mutant immune-competent mouse model. Thus, IAP-JAK-modulatory strategies, like IAP inhibitors, offer promising immunotherapy adjuvants to re-establish the responsiveness of “immunologically-cold” LKB1-mutant tumors to immune checkpoint inhibitors or STING-directed therapies.

## Introduction

Impressive clinical activity of immune system-oriented anti-tumor strategy has led to the rapid rise of immunotherapy as standard care (Aguiar et al., 2017; Hellmann et al., 2018; Hellmann et al., 2019; Robert, 2020; Waldman et al., 2020). However, the diverse primary responsive rate and emerging acquired resistance present a daunting challenge to expand the impact of immunotherapy (Boyero et al., 2020; Sharma et al., 2017; Skoulidis et al., 2018; Skoulidis and Heymach, 2019). In parallel with the immune-targeted effort to search for immune checkpoint molecules and neoantigens, tumor-intrinsic factors have been suggested to modulate tumor immune-responsiveness (Kalbasi and Ribas, 2020; Litchfield et al., 2021; Wellenstein and de Visser, 2018). For example, loss-of-function mutations in β2-microglobulin and Janus kinases (JAK) and amplification of Cyclin D1 have been reported in patients resistant to immunotherapy (Patel et al., 2017; Zaretsky et al., 2016; Zhang et al., 2018). Therefore, understanding how oncogenic drivers determine the intricate tumor immune response may be critical for developing biomarkers and perturbagens to improve immunotherapy efficacy.

Liver kinase B1 (LKB1), also known as Serine/threonine kinase 11, is a tumor suppressor upstream of AMP-activated protein kinases (Marcus and Zhou, 2010; Zhou et al., 2014), and the LKB1-SIK axis has recently been shown to suppress the development of non-small cell lung cancer (Hollstein et al., 2019; Murray et al., 2019). LKB1 inactive mutations (mut) frequently occur in lung adenocarcinoma (LUAD) (Cancer Genome Atlas Research, 2014; Weir et al., 2007). LKB1-mut LUAD tumors gain cellular fitness advantage through mTORC1 activation, and subsequent metabolic rewiring and epigenetic reprogramming (Kottakis et al., 2016). However, LKB1-mut LUAD is considered as “undruggable” due to the loss-of-function mutations found in tumors and the resistance to chemotherapy, targeted therapy and immunotherapy (Calles et al., 2015; El Osta et al., 2019; Skoulidis et al., 2018; Xiao et al., 2016).

In addition to these tumor cell-autonomous hallmarks, the role of LKB1-mut in shaping a suppressive tumor-immune landscape is emerging (Della Corte et al., 2020; Kitajima et al., 2019; Koyama et al., 2016; Skoulidis et al., 2018; Skoulidis and Heymach, 2019). It has been reported that LKB1-mut is a major genetic driver of primary resistance to immune checkpoint inhibitors (Kwack et al., 2020; Pore et al., 2021; Skoulidis et al., 2018). LKB1-mut LUAD have been characterized to exhibit an immune-suppressive phenotype through multiple mechanisms. For example, it has been reported that LKB1-mut LUAD have altered expression of proinflammatory cytokines and immune checkpoint molecules, reprogrammed immune infiltration, and remodeled extracellular matrix (Cristescu et al., 2018; Gao et al., 2010; Gilbert-Ross et al., 2017; Kadara et al., 2017; Koyama et al., 2016; Skoulidis et al., 2015; Skoulidis et al., 2018). It has been demonstrated that LKB1-mut LUAD is associated with repressed expression of stimulator of interferon genes (STING) and corresponding tumor-intrinsic DNA-sensing innate immune response (Della Corte et al., 2020; Kitajima et al., 2019). These LKB1-mut associated “immune cold” features have established LKB1 status as a predictive marker for immunotherapy response. However, the mechanisms underlying the LKB1-mut genotype and immune suppressive phenotype remain to be elucidated. Nevertheless, therapeutic approaches to target LKB1-mut LUAD and reverse its immunosuppression remain absent.

To explore strategies to reverse tumor genotype-dictated immune therapeutic resistance, we took a chemical biology approach to examine the LKB1-mut-created dependency of tumor cells for survival and the underlying molecular basis. Here, we report the identification of IAP (inhibitor of apoptosis protein) antagonists that inhibit the growth of LKB1-mut cells in an immune-dependent manner both *in vitro* and *in vivo*. Functional studies revealed a mechanism by which IAP inhibitors sensitize tumor cells for immune response through the regulation of STING expression and DNA-sensing pathway in LKB1-mut cells. The loss of LKB1 function appears to drive the IAP dependency through the JAK-regulated STING innate immunity pathway in LKB1-mut cells. Modulators of the IAP-JAK regulatory axis, such as the IAP inhibitors, may lead to restored STING pathway function and enhanced sensitivity of LKB1-mut cells to immunotherapeutic assault.

## Results

### Discovery of small molecule immune response sensitizers for LKB1 mutant cells

LKB1-mut LUAD defines a genetic subset of lung cancer with aggressive clinical presentation and therapeutic resistance (Skoulidis et al., 2015; Skoulidis et al., 2018; Xiao et al., 2016). However, the molecular mechanism bridging the LKB1-mut genotype and immune suppression phenotype remains elusive, hampering potential clinical translation. Considering the largely unknown and complex oncogenic signaling underpinning LKB1-mut associated immune suppression, we carried out an unbiased high-throughput immunomodulator phenotypic (HTiP) screen to identify therapeutic vulnerability created by LKB1 mutations, particularly the immune-dependent therapeutic vulnerability.

To examine whether our established HTiP platform (Mo et al., 2019), featuring *in vitro* cancer- and immune-cell co-culture, could recapitulate LKB1-mut associated immune suppression, we tested the response of LUAD cells with LKB1-mut or WT to the immune-cell attack. First, a pair of isogenic cell lines, parental H1792 with LKB1-WT, and the corresponding LKB1 shRNA-knockdown (KD) cells with defined and matched genetic backgrounds, were subjected to the test. Native non-labeled human PBMCs were added to provide the allogenic immune selection pressure. We found that parental H1792 cells with LKB1-WT exhibited high sensitivity to the immune attack with significantly decreased cell viability, whereas the effect was drastically attenuated in isogenic cells with LKB1-KD (Fig. 1A). Similar results were observed in another pair of isogenic H1299 cells (Fig. S1A) and in a panel of patient-derived cancer cells with differential LKB1 status (Fig. 1B). These results recapitulate the LKB1 mutation-associated intrinsic immune resistance.

**Figure 1.**
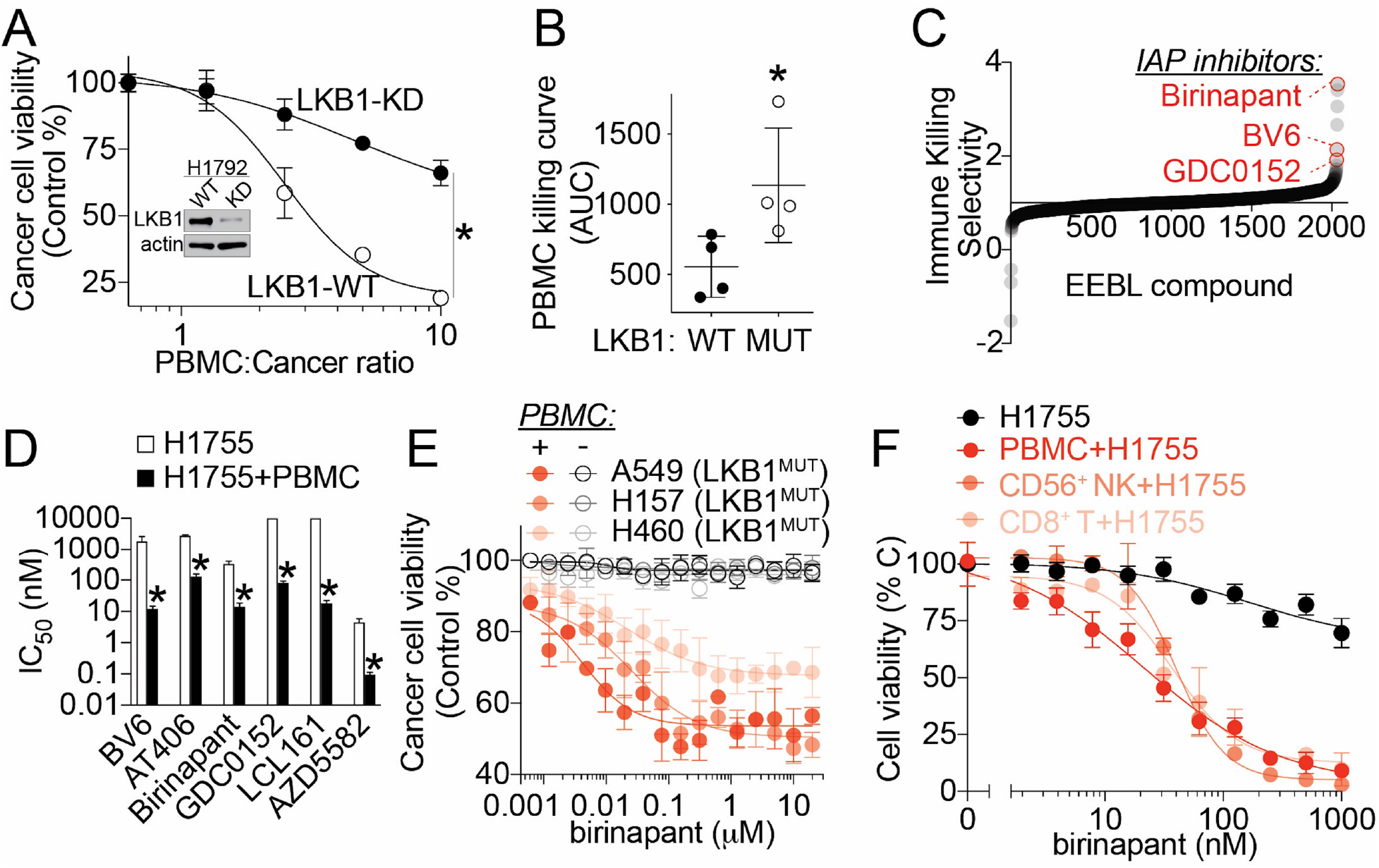
Identification of IAP inhibitors as immune response sensitizers in LKB1-mut lung cancer, related to Fig. S1. **(A)** PBMC dose-response curve showing immune resistance in LKB1-mut cells. Parental LKB1-WT or isogenic LKB1-knockdown (KD) H1792 cells were co-cultured with PBMC at various ratio as indicated. The data are presented as mean±SD from three independent experiments. *p*≤*0.05. **(B)** AUC analysis of PBMC-dose dependent killing curves of a panel of lung cancer cells with LKB1-WT (Calu-1, H1299, H1792 and H292) or MUT (A549, H1792, H23 and H460). Each dot represents one cell line and the line indicate the mean. *p*≤*0.05. **(C)** The waterfall plot showing the selectivity of compounds in immune cell-dependent killing of H1755, a LKB1-mut cell line. The data are presented as the immune killing selectivity from the primary screening. **(D)** Bar graph showing IC_50_s of six structurally diverse IAP inhibitors in H1755 cancer cell alone culture versus co-culture with PBMC. The data are presented as mean±SD from three independent experiments. *p*≤*0.05. **(E)** Dose-response confirmation of birinapant-induced immune-dependent killing in additional LKB1-mut LUAD cell lines as indicated. The data are presented as mean±SD from three independent experiments. **(F)** Dose-response curves of birinapant-induced CD8^+^ T and CD56^+^ NK cell-dependent killing in H1755 cell. The data are presented as mean±SD from three independent experiments.

To identify potential chemical probes to revitalize LKB1-mut cells’ responsiveness to immune selection pressures, we performed an unbiased HTiP screen using a chemogenomic compound library with well-annotated bioactive compounds (Du et al., 2020; Mo et al., 2019; Tang et al., 2021; Yang et al., 2021). This screen revealed that three structurally diverse Inhibitor of Apoptosis Protein (IAP) antagonists, birinapant, BV6 and GDC0152, exhibited immune-dependent killing activity (Fig. 1C and Fig. S1B). To further support the specific impact of targeting IAP for immunomodulation, we tested the effect of three additional IAP inhibitors, AT406, AZD5582 and LCL161. We found that all six IAP inhibitors induced immune cell-dependent selective killing with potency in the nanomolar range (Fig. 1D). Similar effects on immune-dependency of IAP inhibitors were observed in additional LKB1-mut LUAD cell lines, such as A549, H157 and H460 (Fig. 1E). Importantly, birinapant significantly enhanced the response of LKB1-mut LUAD cells to both isolated CD8^+^ T cells and CD56^+^ NK cells (Fig. 1F). These results reveal IAP as a potential immune-dependent vulnerability in LKB1-mut LUAD cells, and targeting IAP may enhance the responsiveness of LKB1-mut tumors to immune-mediated killing. In addition, the immune-dependent anti-tumor activity of IAP inhibitors was more profound towards LKB1-KD cells as compared to the LKB1-WT counterparts (Fig. S1C), indicating potential tumor-intrinsic differentiating factors associated with LKB1 genotype that underly such selectivity.

### Restoration of tumor intrinsic STING expression in LKB1-mut cells by IAP inhibitors

To dissect the downstream tumor-intrinsic differentiating factors that contribute to IAP inhibitor-induced immune-dependent anti-tumor activity towards LKB1-mut cells, we examined several factors, including STING, IL-1*α*, IL-6, G-CSF and PD-L1, that have been reported to shape the LKB1-mut tumor-immune responsiveness (Fig. 2A) (Kitajima et al., 2019; Koyama et al., 2016; Skoulidis et al., 2015; Skoulidis et al., 2018). Given the potent immune-dependent anti-cancer activity of IAP inhibitors, we reasoned that the contributing downstream effectors should also be modulated in a similar immune-dependent manner upon IAP inhibitor treatment. From a focused gene expression profiling of these factors, we found that the STING mRNA level was significantly increased upon birinapant treatment only in the presence of immune cells (Fig. 2A), whereas the mRNA levels of other factors, such as IL-1a, IL-6, GM-CSF and PD-L1, were not significantly changed in an immune-cell dependent manner (Fig. 2A and Fig. S2).

**Figure 2.**
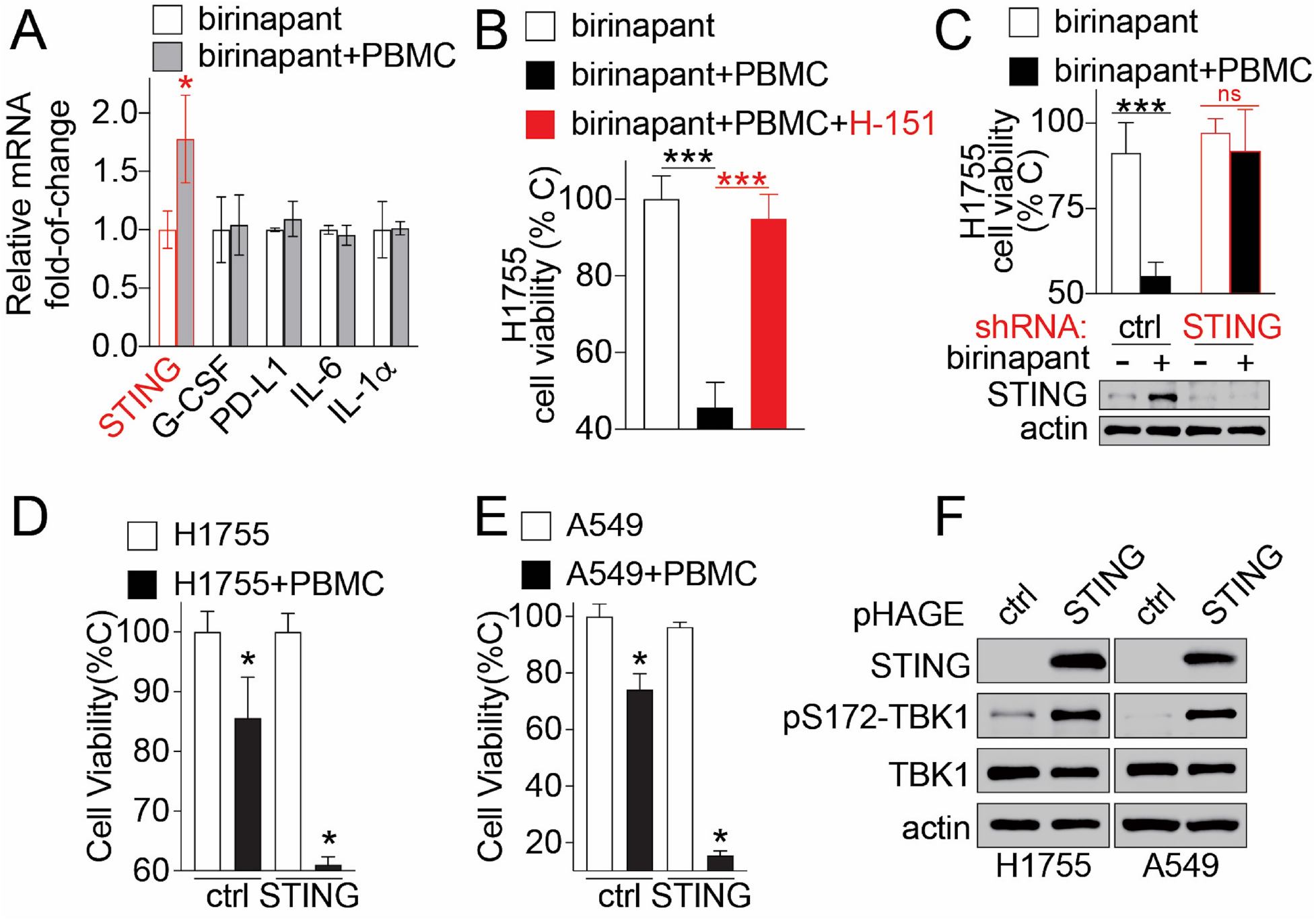
Birinapant sensitizes LKB1-mut cells’ immune response through inducing tumor-intrinsic STING expression, related to Fig. S2-3. **(A)** Bar graph showing birinapant-induced STING gene expression in a immune-dependent manner. H1755 cancer cells were cultured alone or co-cultured with immune cells (E:T=1:1). The change of relative mRNA expression of the selected genes was expressed as fold-of-change upon birinapant (50 nM) treatment over normalized DMSO control. The data are presented as mean±SD from three independent experiments. *p*≤*0.05. **(B)** Bar graph showing of the effect of STING inhibition using H151 on birinapant-induced immune-dependent killing of H1755 cells. The cell viability of H1755 was expressed as percentage of untreated control from H1755 cancer cell alone. H1755 cells were cultured alone or co-cultured with PBMC (E:T=1:1), or treated with birinapant (100 nM) or in combination with H151 (5 μM). The data are presented as mean±SD from three independent experiments. ***p*≤*0.001. **(C)** Bar graph showing the effect of genetic knockdown of STING in H1755 cells on birinapant-induced immune-dependent anti-tumor activity. The cell viability of H1755 was expressed as percentage of untreated control from parental H1755 cancer cell alone. Stable isogenic H1755 cells expressing non-targeting control (ctrl) shRNA or STING-targeting (STING) shRNA were cultured alone or co-cultured with PBMC (E:T=1:1), or treated with birinapant (100 nM) as indicated. Immunoblot (lower) showing birinapant-induced STING expression in control H1755 cells, but not STING knockdown isogenic cells. The data are presented as mean±SD from three independent experiments. ^ns^p>0.05, ***p*≤*0.001. **(D-E)** Bar graph showing STING overexpression-induced increase of immune responsiveness in LKB1-mut cells. Stable isogenic H1755 (D) and A549 (E) cells overexpressing STING were cultured alone or co-cultured with PBMC (E:T=1:1 and 20:1 for H1755 and A549, respectively). The cell viability was expressed as percentage of untreated control from parental cell alone culture. *p*≤*0.05. **(F)** Western blot showing stable isogenic H1755 and A549 cells overexpressing STING.

Suppression of tumor intrinsic STING expression was previously correlated with LKB1 genetic status and the LKB1-mut-associated immune resistance (Fig. S3) (Della Corte et al., 2020; Kitajima et al., 2019). To further test whether STING contributes to IAP inhibitor-enhanced immune responsiveness, we performed chemical and genetic perturbation studies on STING, and examined birinapant-induced immune-dependent anti-tumor activity. Pharmacological loss-of-function study showed that H-151, a STING specific palmitoylation inhibitor (Haag et al., 2018), led to a significant attenuation of birinapant-induced immune-dependent killing activity in LKB1-mut cells (Fig. 2B). Consistently, genetic loss-of-function study via shRNA knockdown of tumor cell intrinsic STING in LKB1-mut cells also significantly blunted IAP inhibitor-induced immune-dependent killing activity (Fig. 2C). In support of this notion, a genetic gain-of-function study showed that overexpression of STING in LKB1-mut cells *per se* did not compromise cancer cells autonomous viability, but significantly sensitized cancer cell’s response to PBMC-mediated immune killing (Fig. 2D-F). Altogether, these results suggested that STING downregulation is a tumor intrinsic and immune-dependent vulnerability in LKB1-mut lung cancer treatment and IAP inhibitors enhance the immune responsiveness of LKB1-mut tumors largely through STING expression restoration.

### IFNγ-JAK1-STAT1 pathway regulates STING expression in LKB1-mut cells

Given that birinapant induced STING expression in an immune-dependent manner, we reasoned that some immune co-factors may contribute and synergize with IAP inhibitors for its immune-dependent anti-tumor activity. To probe the potential immune co-factors, we performed a transcriptome profiling to compare the differential expression genes (DEG) from LKB1-mut tumor cells in the absence or presence of immune cells upon birinapant treatment (Fig. 3A). From the DEG and gene ontology analysis, 1054 immune-dependent upregulated genes induced by birinapant were identified, and these DEGs are enriched in several tumor immune response pathways (Fig. 3B). For example, birinapant activated Type I IFN response signaling (Fig. 3B), a known STING-downstream target pathway (Chen et al., 2016; Zhu et al., 2019), which is suppressed in LKB1-mut LUAD (Fig. S4) (Della Corte et al., 2020; Kitajima et al., 2019). These results further confirmed the immune-dependent activity of birinapant in inducing STING expression.

**Figure 3.**
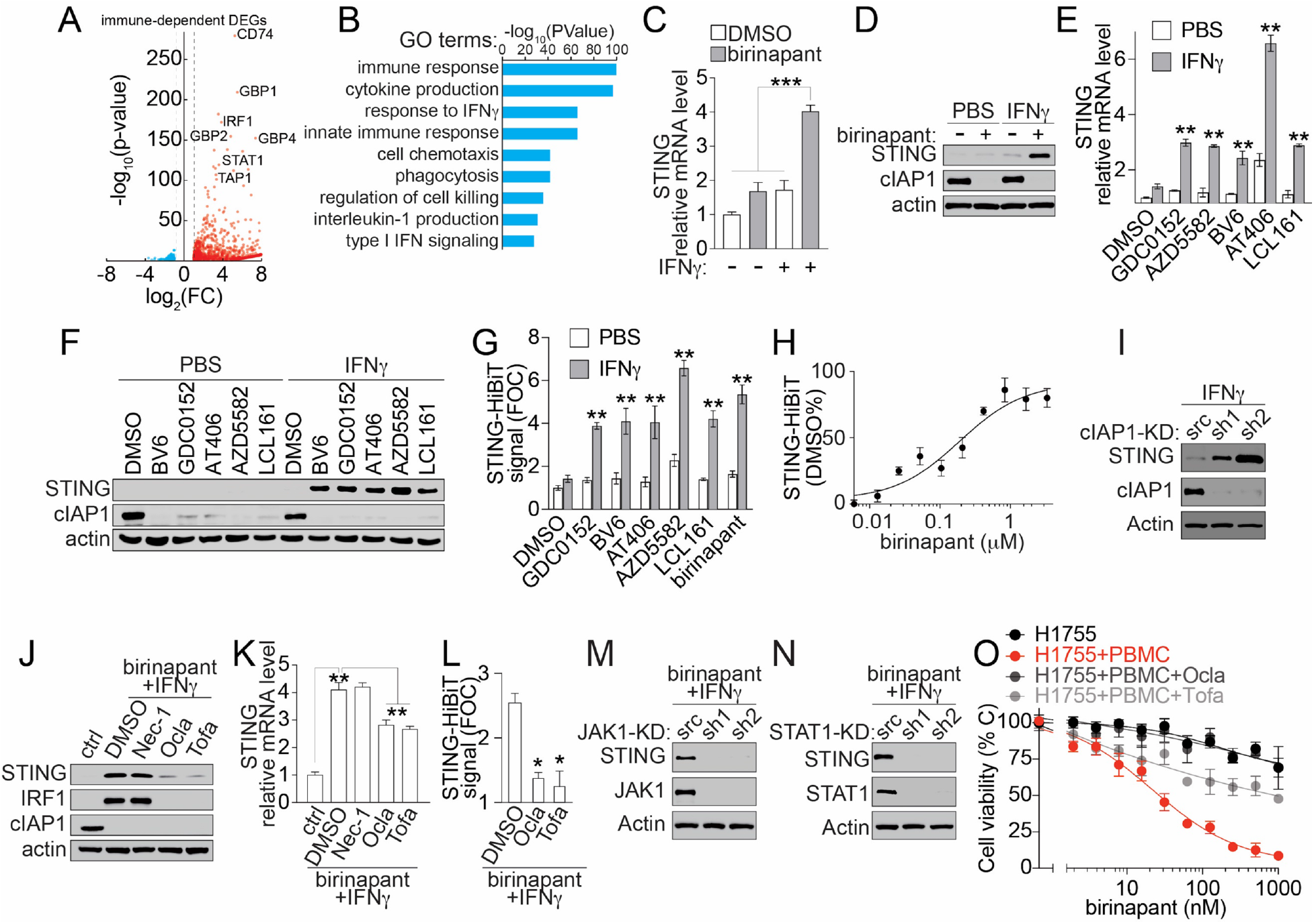
IAP inhibitors synergize with IFNγ to induce STING expression in LKB1-mut cells, related to Fig. S4-6. **(A)** Volcano plot showing the differential expression genes (DEGs) in H1755 cell with immune-dependency. H1755 cells were treated with birinapant (50 nM) in cancer cells alone culture or co-cultured with PBMC (E:T=1:1). The transcriptome of H1755 cells were analyzed by mRNA sequencing. The immune-dependent DEGs were identified using cutoffs of |log_2_(FC)|*≥*1 and adjusted p*≤*0.05. The data are presented as average from three technical replicates. **(B)** Bar graph from gene ontology analysis showing DEGs-associated top-enriched pathways. **(C-D)** STING mRNA (C) and protein (D) expression in A549 cells upon birinapant (500 nM) treatment without (-) or with (+) IFN*γ* (1 ng/mL) for 24 hours. The relative STING mRNA expression was measured using qPCR. The data are presented as mean±SD from three independent experiments. STING protein expression were measured using cell lysate and analyzed using SDS-PAGE with antibodies as indicated. ***p*≤*0.001. **(E-F)** STING mRNA (E) and protein (F) expression in A549 cells upon treatment with IAP inhibitors (500 nM) or in combination with IFN*γ*(1 ng/mL) for 24 hours as indicated. The relative STING mRNA expression was measured using qPCR. The data are presented as mean±SD from three independent experiments. STING protein expression were measured using cell lysate and analyzed using SDS-PAGE with antibodies as indicated. ***p*≤*0.001. **(G)** Bar graph showing STING-HiBiT expression signal in genetically engineered A549 cells treated with various IAP inhibitors (500 nM) or in combination with IFN*γ* (1 ng/mL) for 24 hours as indicated. The STING-HiBiT signal was expressed as the fold-of-change over the DMSO control and presented as mean±SD from three independent experiments. ***p*≤*0.001. **(H)** Dose-response curve of birinapant-induced STING-HiBiT expression in genetically engineered A549 cells in the presence of 1 ng/mL IFN*γ*. The STING-HiBiT signal was expressed as the percentage of DMSO control and presented as mean±SD from three independent experiments. **(I)** Immunoblot showing STING expression in isogenic cIAP1 knockdown A549 cells. A549 cells were treated with 1 ng/mL IFNg for 24 hours, and cell lysates from isogenic A549 cells stably expressing control scramble (scr), or cIAP1 targeting shRNAs (sh1 and sh2), were analyzed by SDS-PAGE with antibodies as indicated. **(J-K)** STING protein (J) and mRNA (K) expression in A549 cells treated with birinapant (500 nM) and IFN*γ* (1 ng/mL) in combination with JAK inhibitors, oclacitinib (Ocla, 10 μM) and tofacitinib (Tofa, 10 μM), or RIPK inhibitor, necrostatin-1 (Nec-1, 10 μM), as indicated. STING protein expression were measured using cell lysates and analyzed by SDS-PAGE with antibodies as indicated. The relative STING mRNA expression was measured using qPCR. The data are presented as mean±SD from three independent experiments. **p*≤*0.01. **(L)** Bar graph showing STING-HiBiT signal in genetically engineered A549 cells treated with birinapant (500 nM) and IFN*γ* (1 ng/mL) in combination with JAK inhibitors, oclacitinib (Ocla, 10 μM) and tofacitinib (Tofa, 10 μM), as indicated. The STING-HiBiT signal was expressed as the fold-of-change over the DMSO control and presented as mean±SD from three independent experiments. *p*≤*0.001. **(M-N)** Immunoblot showing STING expression in isogenic JAK1 (M) or STAT1 (N) knockdown (KD) A549 cells. A549 cells were treated with birinapant (500 nM) and 1 ng/mL IFN*γ*for 24 hours, and cell lysates from isogenic A549 cells stably expressing control scramble (scr), or JAK1 or STAT1 targeting shRNAs (sh1 and sh2), were analyzed by SDS-PAGE with antibodies as indicated. **(O)** Dose-repsonse curve of birinapant-induced growth inhibition of H1755 cells in cancer cell alone culture or co-culture with immune cells in combination with JAK inhibitors, oclacitinib (Ocla, 10 μM) and tofacitinib (Tofa, 10 μM), as indicated. The cell viability of H1755 cells were expressed as percentage of DMSO control and presented as mean±SD from three independent experiments.

In addition, interferon-gamma (IFN*γ*) response pathway was activated by birinapant in a similar immune-dependent manner (Fig. 3A-B), while IFN*γ* pathway genes were significantly downregulated in LKB1-mut LUAD samples (Fig. S4). Meanwhile, we and others have demonstrated that IAP inhibitors induce IFN*γ* production from immune cells (Dougan et al., 2010; Mo et al., 2019). Altogether, we reasoned that IFN*γ* might be an immune co-factor that synergizes with an IAP inhibitor for its immune-dependent anti-tumor and STING-induction activity.

To test this hypothesis, we next queried the contribution of IFN*γ* to birinapant-induced STING restoration. In the culture of cancer cells alone, birinapant synergized with supplemented IFN*γ* to induce STING mRNA and protein expression in LKB1-mut cancer cells (Fig. 3C-D and Fig. S5A-D). In addition, all six IAP inhibitors exhibited similar synergistic effects with IFN*γ*to induce STING mRNA and protein expression (Fig. 3E-F and Fig. S5E-F). Such synergistic effect between IAP inhibitors and IFN*γ* was confirmed at the protein level using a quantitative STING-HiBiT assay (Schwinn et al., 2018), a genetically engineered A549 cell-based reporter system for monitoring endogenous STING expression (Fig. 3G and Fig. S6). Using the STING-HiBiT assay, we found that birinapant induced a dose-dependent increase of STING protein in LKB1-mut A549 cells with EC_50_∼0.2 μM in the presence of IFN*γ* (Fig. 3H). The on-target effect of IAP inhibitor-induced STING expression was further confirmed by the genetic loss-of-function perturbation of cIAP1 (Fig. 3I). These results suggest that IFN*γ* is an immune co-factor that synergize with IAP inhibitors to restore STING expression through targeting cIAP1.

Given that such synergistic effects were also observed at the STING mRNA level (Fig. 3 and Fig. S5), we hypothesized that certain transcriptional machinery induced by IAP inhibitor and IFN*γ*combination treatment might be involved. To further decipher the downstream transcriptional machineries that were activated for the restoration of STING expression by IAP inhibitor and IFN*γ* combination, we examined RIPK and JAK-STAT signaling, two reported effectors downstream of IFN*γ* (Hao and Tang, 2018; Tanzer et al., 2017). We found that IAP inhibitor-induced STING restoration was significantly mitigated upon treatment with JAK inhibitors, but not RIPK inhibitors, at both protein and mRNA levels (Fig. 3J-K and Fig. S5G-H). The results were confirmed with STING-HiBiT engineered A549 cells (Fig. 3L). In support of this notion, shRNA knockdown of JAK1 and STAT1 significantly abolished the synergistic effect of IAP inhibitor and IFN*γ* on restoring STING expression in LKB1-mut cells (Fig. 3M-N). Consistently, JAK inhibitors abolished the responsiveness of LKB1-mut tumor cells to IAP inhibitor-induced immune-dependent anti-tumor activity (Fig. 3O). These results demonstrate that IAP inhibitor-induced STING restoration and immune-dependent anti-tumor response in LKB1-mut cells is through the activation and sensitization of the tumor intrinsic IFN*γ*-JAK-STAT pathway.

### Reactivation of STING-mediated DNA-sensing signaling by birinapant in LKB1-mut cells

Suppression of STING expression in LKB1-mut LUAD was demonstrated to lead to immune suppression via impaired cGAS-STING DNA-sensing innate immune response signaling and subsequent inhibition of the downstream TBK1-IRF3 pathway and the expression of IRF3 target cytokine and chemokine genes (Della Corte et al., 2020; Kitajima et al., 2019). Given that IAP inhibitors restore STING expression, we next queried whether IAP inhibitors can restore the STING downstream TBK1-IRF3 signaling and target gene expression.

At the basal level without supplementation of exogenous dsDNA or cyclic dinucleotide, birinapant synergized with IFN*γ* to activate STING downstream signaling with significantly increased levels of p-TBK1 and p-IRF3 (Fig. 4A) and expression of target cytokine and chemokine genes, such as IFN*β*, CXCL10 and CCL5 (Fig. 4B-D). These results suggest that IAP inhibition not only restores STING expression, but also reactivates STING-mediated DNA-sensing signaling in response to cytosolic DNA, which could be induced by an IAP inhibitor or IFN*γ* treatment (Fig. S7) or may arise from LKB1-mut tumor intrinsic genome instability (Gupta et al., 2015).

**Figure 4.**
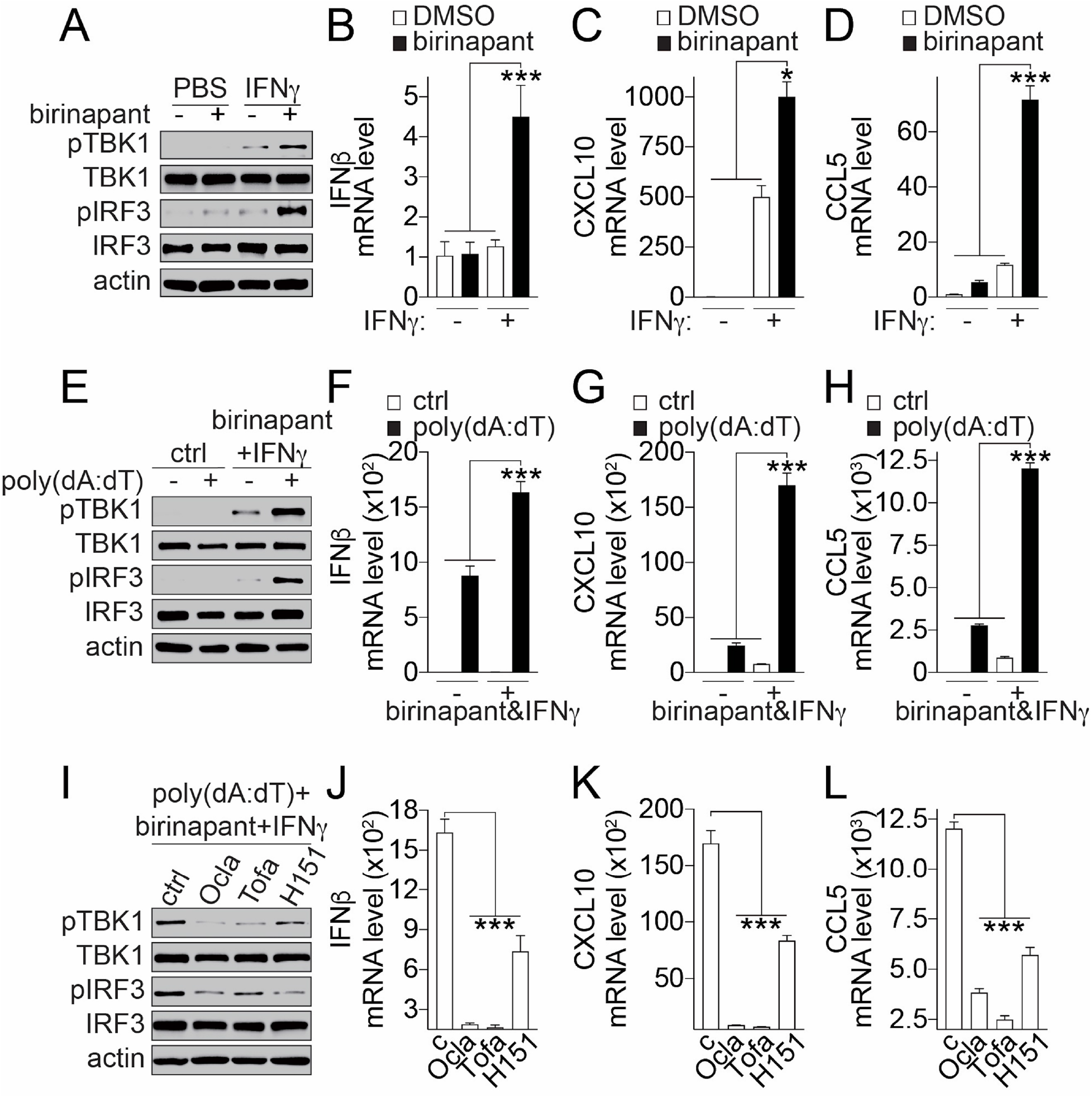
IAP inhibitors synergize with IFNγ to induce STING-mediated DNA sensing pathway activation in LKB1-mut cells, related to Fig. S7-8. **(A)** Immunoblot showing birinapant and IFN*γ* combination-induced activation of TBK1 and IRF3. A549 cells were treated with birinapant (500 nM) and/or IFN*γ* (1 ng/mL) for 24 hours as indicated. Cell lysate were analyzed by SDS-PAGE and western blot with antibodies as indicated. **(B-D)** Bar graphs showing birinapant and IFN*γ* combination-induced expression of IFN*β* (B), CXCL10 (C) and CCL5 (D). A549 cells were treated with birinapant (500 nM) and/or IFN*γ* (1 ng/mL) for 24 hours as indicated. Gene expression were analyzed by qPCR and presented as mean±SD from three independent experiments. *p*≤*0.05, *** p*≤*0.001. **(E)** Immunoblot showing poly(dA:dT)-induced TBK1 and IRF3 activation. A549 cells were treated with poly(dA:dT) (1 μg/mL) for 4 hours in the presence or absence of 24-hour pre-treatment with birinapant (500 nM) and IFN*γ* (1 ng/mL) combination as indicated. Cell lysate were analyzed by SDS-PAGE and western blot with antibodies as indicated. **(F-H)** Bar graphs showing poly(dA:dT)-induced expression of IFN*β* (F), CXCL10 (G) and CCL5 (H). A549 cells were treated with poly(dA:dT) (1 μg/mL) for 4 hours in the presence or absence of 24-hour pre-treatment with birinapant (500 nM) and IFN*γ*(1 ng/mL) combination as indicated. Gene expression were analyzed by qPCR and presented as mean±SD from three independent experiments. *** p*≤*0.001. **(I)** Immunoblot showing JAK- and STING-dependency of birinapant-induced reactivation of TBK1 and IRF3 phosphorylation. A549 cells were treated with poly(dA:dT) (1 μg/mL) for 4 hours in the presence of 24-hour pre-treatment of birinapant (500 nM) plus IFN*γ* (1 ng/mL) in combination with JAK inhibitors, oclacitinib (Ocla, 10 μM) and tofacitinib (Tofa, 10 μM), or STING antagonist, H151 (5 μM), as indicated. Cell lysate were analyzed by SDS-PAGE and western blot with antibodies as indicated. **(J-L)** Bar graphs showing JAK- and STING-dependency of birinapant-induced expression of IFN*β* (J), CXCL10 (K) and CCL5 (L). A549 cells were treated with poly(dA:dT) (1 μg/mL) for 4 hours in the presence of 24-hour pre-treatment of birinapant (500 nM) plus IFN*γ*(1 ng/mL) in combination with JAK inhibitors, oclacitinib (Ocla, 10 μM) and tofacitinib (Tofa, 10 μM), or STING antagonist, H151 (5 μM), as indicated. Gene expression were analyzed by qPCR and presented as mean±SD from three independent experiments. *** p*≤*0.001.

Upon further STING activation using poly(dA:dT), an exogenous dsDNA, we found that birinapant and IFN*γ* combination treatment significantly augmented dsDNA-induced TBK1 and IRF3 phosphorylation (Fig. 4E) and expression of IFN*β*, CXCL10 and CCL5 (Fig. 4F-H). The augmentation effect of the TBK1-IRF3 signaling by the birinapant and IFN*γ* combination was not limited to A549 cells. Similar effects were observed in additional LKB1-mut cells at basal level (Fig. S8A-D) or with dsDNA stimulation (Fig. S8E-H). In addition, the birinapant and IFN*γ* combination treatment also significantly enhanced LKB1-mut cells’ response to ADU-S100, a synthetic cyclic dinucleotide STING agonist (Corrales et al., 2015), in terms of TBK1 and IRF3 phosphorylation (Fig. S8I). Further, we found that a JAK or STING inhibitor abolished TBK1 and IRF3 phosphorylation (Fig. 4I) and the expression of IFN*β*, CXCL10 and CCL5 (Fig. 4J-L) induced by dsDNA, indicating the underlying JAK- and STING-dependency. Altogether, these results suggest that LKB1-mut cells with STING downregulation have impaired DNA-sensing signaling, which can be rescued by IAP inhibition through the IFN*γ*-JAK-STAT pathway-mediated STING expression.

### Birinapant induces STING-mediated apoptosis of LKB1-mut cells and chemotaxis of immune cells *in vitro*

To explore thefunctional consequence of IAP inhibitor-induced STING expression and activation, we examined the fate of LKB1-mut cells and immune cell infiltration *in vitro*. STING activation has been shown to induce apoptosis in a context-specific manner (Gaidt et al., 2017; Larkin et al., 2017; Tang et al., 2016; White et al., 2014). Given that IAP inhibitors enhance STING expression and activation in combination with IFN*γ*, we queried whether the combination of an IAP inhibitor with IFN*γ* affect apoptosis of LKB1-mut cells. We found that birinapant alone did not lead to significant cell death, while the combination of birinapant with IFN*γ* induced significant caspase3/7 activation in LKB1-mut A549 cells (Fig. 5A-B). Similar effects of the birinapant and IFN*γ* combination on caspase3/7 activation were also observed in additional LKB1-mut cells (Fig. S9A-B). In addition, such a combination effect was significantly attenuated upon STING inhibition by H151 (Fig. 5C-D). These results reveal that IAP inhibition can synergize with IFN*γ* to induce apoptosis of LKB1-mut tumors in a STING-dependent manner.

**Figure 5.**
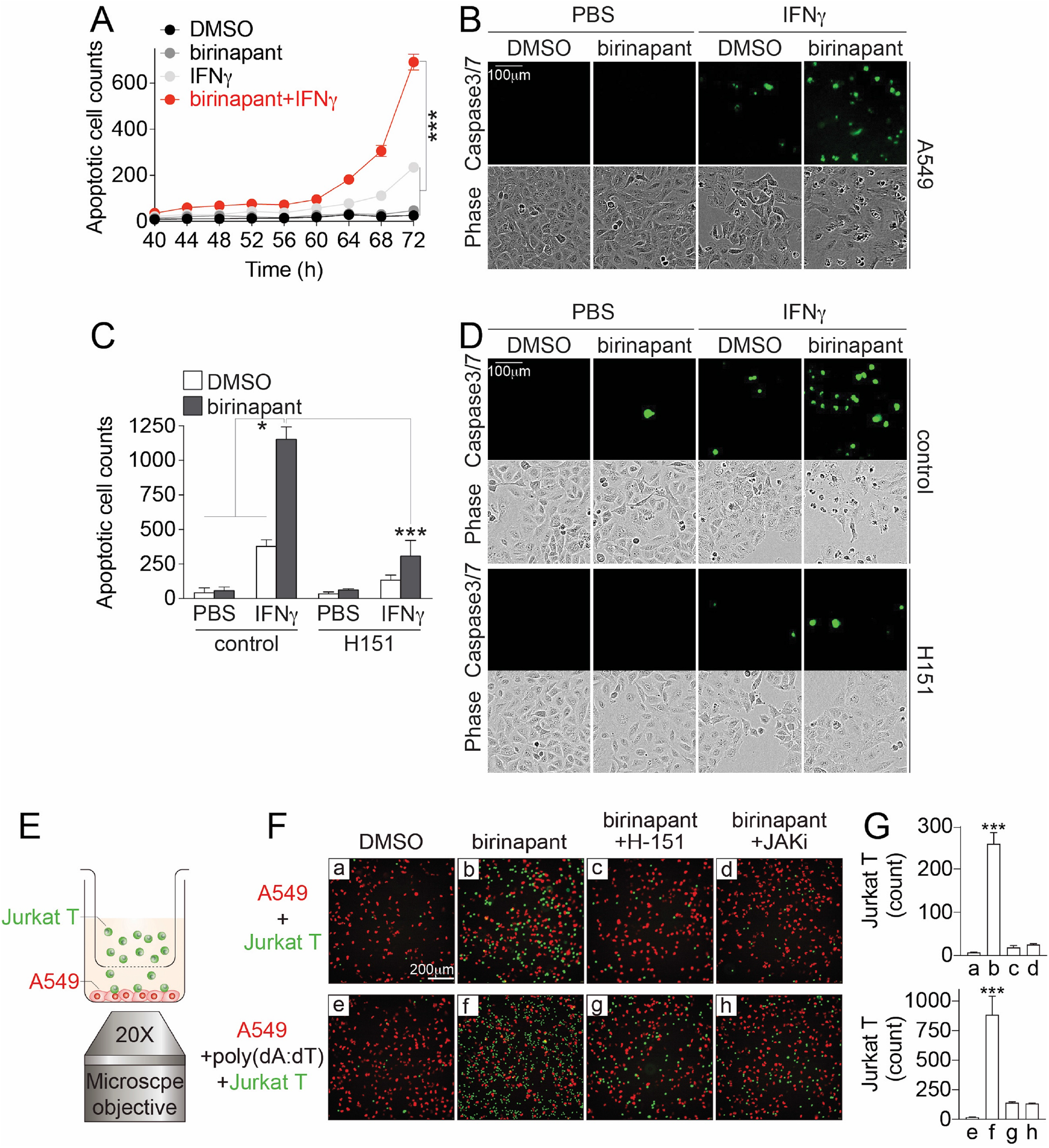
Birinapant induces STING-mediated apoptosis of LKB1-mut cancer cells and chemotaxis of immune cells *in vitro*, related to Fig. S9. **(A)** Time-dependent curve of birinapant-induced cell apoptosis. A549 cells were treated with birinapant (500 nM), IFN*γ* (1 ng/mL) or in combination as indicated. Cell apoptosis was measured and monitored in real-time using DEVD-based fluorogenic caspase-3/7 apoptosis reporter assay up to 72 hours. The data were expressed as the count of green fluorescence positive cells and presented as mean±SD from three independent experiments. *** p*≤*0.001. **(B)** Representative images showing birinapant-induced cell apoptosis. A549 cells were treated with birinapant (500 nM), IFN*γ* (1 ng/mL) or in combination as indicated. Green fluorescence (caspase-3/7 apoptosis reporter) and phase-contrast images were acquired using IncuCyte. **(C)** Bar graph showing birinapant-induced STING-mediated cancer cell apoptosis. A549 cells were treated with birinapant (500 nM), IFN*γ* (1 ng/mL), H151 (5 μM) or in combination as indicated for 72 hours. Cell apoptosis was measured at the endpoint using DEVD-based fluorogenic caspase-3/7 apoptosis reporter assay. The data were expressed as the count of green fluorescence positive cells and presented as mean±SD from three independent experiments. * p*≤*0.05, *** p*≤*0.001. **(D)** Representative images showing birinapant-induced STING-mediated cancer cell apoptosis. A549 cells were treated with birinapant (500 nM), IFN*γ* (1 ng/mL), H151 (5 μM) or in combination as indicated for 72 hours. Green fluorescence (caspase-3/7 apoptosis reporter) and phase-contrast images were acquired using IncuCyte. **(E)** Schematic illustration of transwell assays for measuring immune cell infiltration *in vitro*. **(F)** Representative fluorescence images showing birinapant-induced Jurkat T cells migration. A549 cells and Jurkat T cells were co-cultured in transwell as shown in (E) and were treated with birinapant (100 nM), poly(dA:dT) (1 μg/mL), or in combination with STING antagonist, H151 (5 μM), or JAK inhibitor (JAKi), tofacitinib (10 μM), for 48 hours as indicated. Fluorescence images of A549 cells (red) and infiltrated Jurkat T cells (green) were acquired at the endpoint using ImageXpress Micro high-content imaging system. **(G)** Bar graph showing the quantification of infiltrated Jurkat T cells in transwell-based migration assays. The data were expressed as the count of green fluorescence positive Jurkat T cells and presented as mean±SD from three independent experiments. * p*≤*0.05, *** p*≤*0.001.

Given that STING-mediated CXCL10 and CCL5 chemokines are involved in immune cell chemotaxis (Chan et al., 2012; Tokunaga et al., 2018), we next examined the potential functional consequence of IAP inhibitor-induced STING expression on chemotaxis of immune cell in a 2D transwell assay. As shown in Fig. 5E, the birinapant treatment significantly promoted the migration of Jurkat T cells from the upper chamber to the cancer cell culture in the bottom chamber (Fig. 5F-G). Similar effects of birinapant-induced immune cell migration were observed for CD56^+^ NK92-MI cells or with additional dsDNA stimulation (Fig. 5F-G and Fig. S9C-E). However, a JAK inhibitor, or a STING antagonist, abolished the birinapant-induced immune cell migration (Fig. 5F-G and Fig S9C-E). These results suggest that IAP inhibitor-induced STING expression and reactivation can augment immune cell migration *in vitro*.

### Birinapant restores STING expression and exhibits immune-dependent anti-tumor activity *in vivo*

To determine the functional consequence of IAP inhibitor-induced STING expression and reactivation in an *in vivo* setting we examined the anti-tumor activity of IAP inhibitors using our recently developed LKB1-mut mouse model (Gilbert-Ross et al., 2017; Jin et al., 2021). First, we found that WRJ388, a mouse tumor-derived cancer cell line isolated from our *Kras^G12D^/Lkb1^-/-^/p53^WT^* GEMM (Gilbert-Ross et al., 2017), has downregulated STING expression as compared to KW634 cells (Ji et al., 2007) (*Kras^G12D^/Lkb1^WT^/p53^-/-^*) (Fig. S10A-B). Moreover, IAP inhibitors in combination with mouse IFN*γ* significantly induced STING expression (Fig. S10C-D) and CCL5 expression (Fig. S10E) in *Lkb1*-mut WRJ388 cells. These results suggest that WRJ388 cells can recapitulate not only the LKB1-mut and STING-downregulation genotype-phenotype relationship as observed in human LUAD patients, but also the IAP dependency as shown in human LKB1-mut cells.

Using the syngeneic allograft model with WRJ388 cells, we next assessed the anti-tumor activity of birinapant *in vivo* in LKB1-mut background. Murine WRJ388 cells were subcutaneously injected into the syngeneic immunocompetent mice followed by birinapant treatment (Fig. 6A). We found that treatment with birinapant led to a ∼42% tumor volume shrinkage from ∼300 to ∼175 mm^3^, which is ∼4.4-fold less than the vehicle control at the endpoint, showing a significant anti-tumor effect (Fig. 6B). By contrast, no significant tumor shrinkage was observed with the birinapant treatment in the immune-deficient nude mice (Fig. 6C). Thus, birinapant exhibits immune-dependent anti-tumor activity *in vivo*.

**Figure 6.**
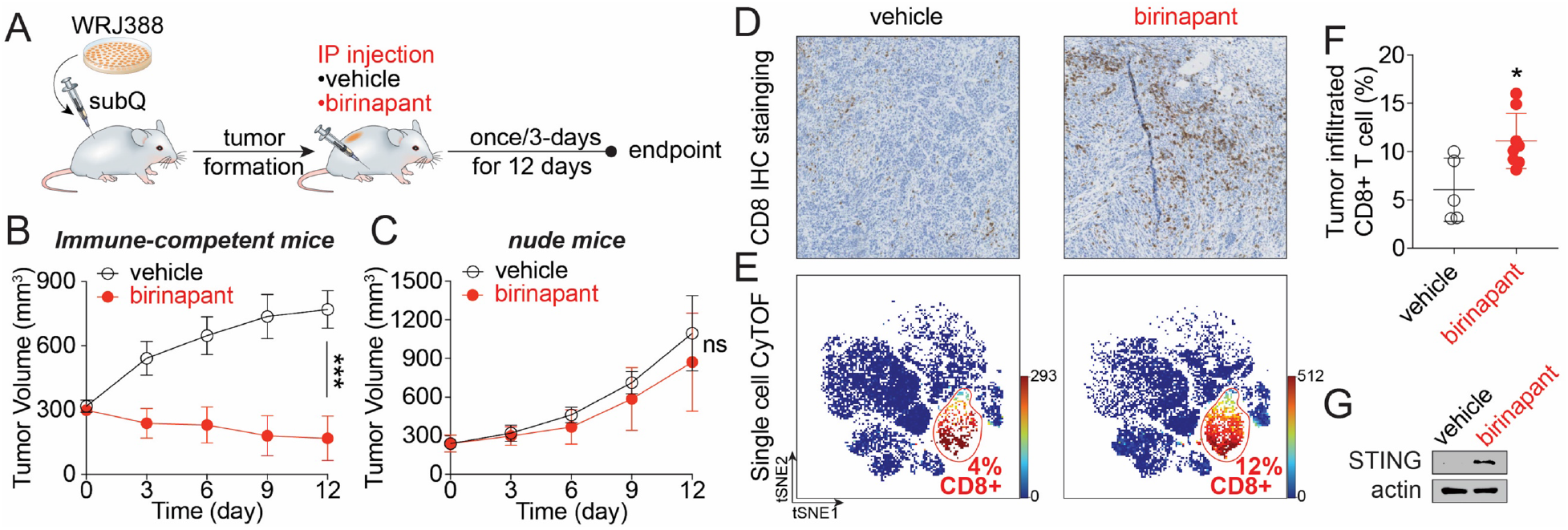
Birinapant exhibits immune-dependent anti-tumor activity *in vivo* in a LKB1-mut syngeneic allograft mouse model, related to Fig. S10. **(A)** Schematics of mouse study design. **(B-C)** Time course of tumor volume change in immune-competent mice (B, n=6 per group) and immune-deficient nude mouse (C, n=5 per group) treated with vehicle control or birinapant (10 mg/kg) as indicated. The data are presented as mean±SD of the entire experimental cohort. ^ns^p>0.05, *** p*≤*0.001. **(D)** Representative images from immunohistochemistry (IHC) staining showing 2-doses birinapant-induced increase of tumor infiltrated CD8^+^ T cells *in vivo*. **(E)** tSNE plots from single cell mass cytometry (CyTOF) showing increased tumor infiltrated CD8+ T cells upon 2-doses birinapant treatment *in vivo*. **(F)** Quantification of tumor infiltrated CD8^+^ T cells from single cell CyTOF profiling. The data are expressed as the percentage of CD8^+^ T cells in live cell population. Each data point represents individual samples from immune-competent mice. *p*≤*0.05 **(G)** Immunoblot showing birinapant-induced STING expression *in vivo*. Tumor sample harvested from the immune-competent mice at the endpoint were analyzed by SDS-PAGE and western blot with indicated antibodies.

Tumor samples from the immune-competent mice were then analyzed to determine the effect of birinapant on the tumor immune microenvironment. The number of tumor infiltrated CD8^+^ T cells was significantly increased upon birinapant treatment of as shown by immunohistochemistry staining (Fig. 6E). Such increase in tumor infiltrated CD8^+^ T cells was confirmed using unbiased single-cell mass cytometry profiling (Fig. 6F-G). Significantly, birinapant treatment restored STING expression *in vivo*, supporting our results from *in vitro* studies (Fig. 6G). Altogether, these results demonstrate that birinapant restores STING expression in LKB1-mut tumors to exhibit enhanced cytotoxic T cell infiltration and immune-dependent anti-tumor activity in a syngeneic mouse model.

### Identification of LKB1-cIAP1-JAK1 trimolecular interactions for the JAK1-STING regulation

The molecular and cellular functional studies *in vitro* and *in vivo* above revealed the IAP-STING regulatory axis as a potential immune-dependent vulnerability in LKB1-mut cells. The IAP inhibitor action appears to be associated with the IFN*γ*-JAK-STAT pathway because IAP inhibitors show a synergistic effect with IFN*γ*for their roles in restoring STING expression and activation. However, the molecular mechanisms that determine the IAP-dependency in the context of LKB1-mut state of tumor cells and their resistance to immune attacks remain to be established.

To understand the potential molecular mechanisms, we performed a binary protein-protein interaction (PPI) profiling among LKB1, cIAP1 and IFN*γ*-JAK-STAT pathway proteins. Using a robust TR-FRET PPI mapping assay (Ivanov et al., 2017; Ivanov et al., 2018; Li et al., 2017; Mo et al., 2017), we found that cIAP1 strongly interacted with LKB1 and JAK1, respectively (Fig. 7A and Fig. S11A), suggesting an intricate interplay among LKB1, cIAP1, and JAK1. The binary LKB1-cIAP1 and cIAP1-JAK1 PPIs were further confirmed by GST-beads affinity pulldown of the overexpressed proteins (Fig. 7B-C) and co-immunoprecipitation of endogenous proteins (Fig. 7D-E). These results support the physical interaction of LKB1 and JAK1 with cIAP1 under physiologically relevant conditions.

**Figure 7.**
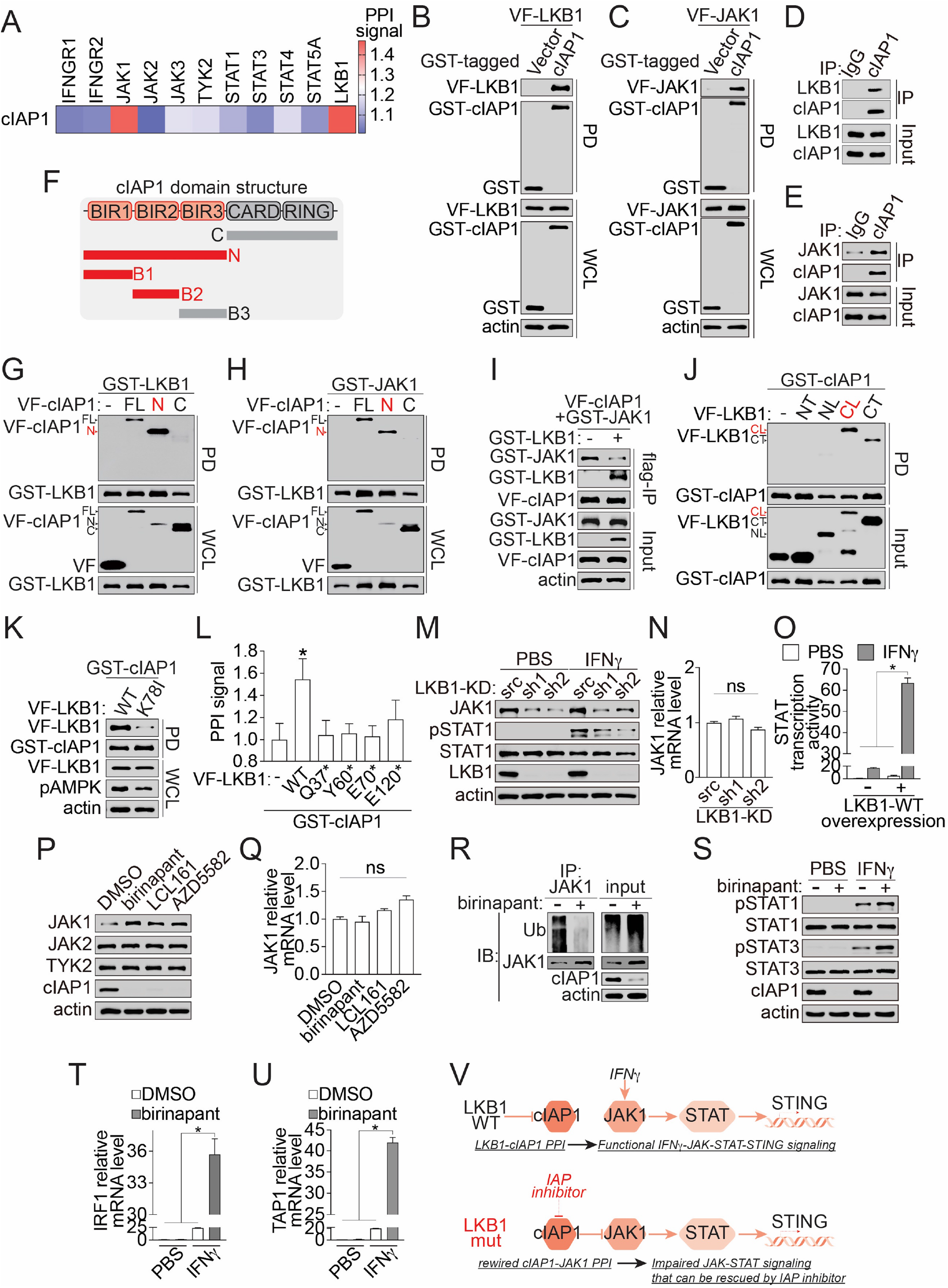
Identification of the LKB1-cIAP1-JAK1 trimolecular complex in shaping IAP- and STING-dependency in LKB1-mut cells, related to Fig. S11. **(A)** Heatmap showing PPI signal between cIAP1 and LKB1, or IFN*γ*-JAK-STAT pathway proteins. PPI signal were measured using cell lysate from HEK293T cells co-expressing GST-cIAP1 and Venus-flag (VF)-tagged test proteins. The PPI signals were expressed as the fold-of-change over the corresponding empty vector controls and presented as the average of three technical replicates from the primary screen. **(B-C)** Immunoblot showing GST-beads affinity pulldown confirmation of LKB1-cIAP1 (B) and cIAP1-JAK1 (C) PPI. Cell lysate from HEK293T cells co-expressing GST-cIAP1 and Venus-flag (VF)-tagged LKB1 or JAK1 were subjected to the GST-pulldown as indicated. The pulldown complex and whole cell lysate (WCL) were analyzed by SDS-PAGE and western blot with antibodies as indicated. **(D-E)** Endogenous interaction of LKB1-cIAP1 (D) and cIAP1-JAK1 (E) with the co-IP assay. The cIAP1 PPI complex was immunoprecipitated from H1299 (LKB1-WT) and A549 (LKB1-mut) cell lysate for confirming LKB1-cIAP1 (D) and cIAP1-JAK1 (E) PPI, respectively. The co-IP complex and whole cell lysate input sample was analyzed by immunoblotting as indicated. **(F)** Schematic illustration of the design of cIAP1 domain truncations **(G-H)** Immunoblot showing mapping of LKB1- (G) and JAK1- (H) binding domain on cIAP1. Cell lysate from HEK293T cells co-expressing GST-LKB1 or JAK1 with and Venus-flag (VF)-tagged cIAP1 full-length (FL), N-terminal truncation (N) or C-terminal truncation (C) were subjected to the GST-pulldown as indicated. The pulldown complex and whole cell lysate (WCL) were analyzed by SDS-PAGE and western blot with antibodies as indicated. **(I)** Immunoblot showing competitive binding between LKB1 and JAK1 with cIAP1. Cell lysate from HEK293T cells co-expressing GST-JAK1 and Venus-flag (VF)-tagged cIAP1 with (+) or without (-) additional GST-LKB1 were subjected to the flag-beads immunoprecipitation (flag-IP) as indicated. The VF-cIAP1 PPI complex from flag-IP and whole cell lysate input were analyzed by SDS-PAGE and western blot with antibodies as indicated. **(J)** Immunoblot showing mapping of cIAP1-binding domain on LKB1. Cell lysate from HEK293T cells co-expressing GST-cIAP1 and Venus-flag (VF)-tagged LKB1 N-terminal truncation (NT, 1-42 aa), kinase domain N-lobe truncation (NL, 43-131aa), kinase domain C-lobe truncation (CL, 132-347aa), and C-terminal truncation (CT, 348-433aa) were subjected to the GST-pulldown as indicated. The pulldown complex and whole cell lysate (WCL) were analyzed by SDS-PAGE and western blot with antibodies as indicated. **(K)** Immunoblot showing the LKB1 kinase-dependency of LKB1-cIAP1 PPI. Cell lysate from HEK293T cells co-expressing GST-cIAP1 and Venus-flag (VF)-tagged LKB1 WT or K78I kinase-dead mutant were subjected to the GST-pulldown as indicated. Before collecting lysate, cells were serum-starved for overnight. The pulldown complex and whole cell lysate (WCL) were analyzed by SDS-PAGE and western blot with antibodies as indicated. **(L)** Bar graph showing PPI signal between cIAP1 and LKB1 WT or patient-derivedmutants. Cell lysate from HEK293T cells co-expressing GST-cIAP1 and Venus-flag (VF)-tagged LKB1 WT or mutants as indicated were subjected to the TR-FRET assay. The PPI signal was expressed as fold-of-change of the TR-FRET signal over the empty vector control (-) and presented as mean±SD from three independent experiments. *p*≤*0.05. **(M-N)** JAK1 protein (M) and mRNA (N) expression in isogenic LKB1-knockdown (KD) H1299 cells. H1299 cells were treated with 1 ng/mL IFN*γ* or PBS control for 24 hours. JAK1 protein level was measured using cell lysates from isogenic H1299 cells stably expressing control scramble (src), or LKB1 targeting shRNAs (sh1 and sh2), and were analyzed by SDS-PAGE with antibodies as indicated. The relative JAK1 mRNA expression was measured using qPCR. The data are presented as mean±SD from three independent experiments. ^ns^p>0.05. **(O)** Bar graph showing LKB1 WT overexpression-induced STAT-driven ISRE transcriptional activation. HEK293T cells were transfected with ISRE-luc reporter plasmid with (+) or without (-) LKB1-WT overexpression, and were treated with 1 ng/mL IFN*γ* or PBS control as indicated for 24 hours. The ISRE-luc reporter activity were measured by Dual-Glo luciferase kits and are presented as mean±SD from three independent experiments. *p*≤*0.05. **(P-Q)** JAK1 protein (P) and mRNA (Q) expression upon IAP inhibitor treatment in LKB1-mut cells. A549 cells were treated with IAP inhibitors (500 nM) as indicated for 24 hours. JAK1 protein level was measured using cell lysates and were analyzed by SDS-PAGE with antibodies as indicated. The relative JAK1 mRNA expression was measured using qPCR. The data are expressed as fold-of-change over the DMSO control (FOC) and presented as mean±SD from three independent experiments. ^ns^p>0.05 and FOC<1.5. **(R)** Representative immunoblot showing birinapant-induced decrease of JAK1 ubiquitination in LKB1-mut cells. Cell lysate from A549 cells with 18-hour birinapant (500 nM) followed by 6-hour MG132 (20 μM) treatment were subjected to endogenous JAK1 immunoprecipitation (IP). Immunoprecipitated complex and whole cell lysate input was analyzed by SDS-PAGE and western blot with antibodies indicated. **(S)** Immunoblot showing birinapant-induced sensitization of IFN*γ*-mediated STAT1/3 activation in LKB1-mut cells. A549 cells were treated with IFN*γ* (1 ng/mL) or PBS control for 1 hour with (+) or without 24-hour pretreatment of birinapant (500 nM). Cell lysate were analyzed by SDS-PAGE and western blot with antibodies as indicated. **(T-U)** Bar graphs showing birinapant-induced sensitization of IFN*γ*-mediated IRF1 (S) and TAP1 (T) mRNA expression. A549 cells were treated with birinapant (500 nM), IFN*γ* (1 ng/mL), or in combination as indicated for 24 hours. The relative IRF1 and TAP1 mRNA expression was measured using qPCR. The data are expressed as fold-of-change over the PBS control (FOC) and presented as mean±SD from three independent experiments. *p*≤*0.05. **(V)** A working model showing the effect of the LKB1 status –on the interact of cIAP1 with JAK1 in regulating STING expression via IFN*γ*-JAK-STAT pathway modulation.

To understand the mode of interaction, we examined the structural determinants of cIAP1 in mediating its interaction with LKB1 and JAK1. cIAP1 domain-based truncation studies were carried out, which showed that both LKB1 and JAK1 were associated with the same domain of cIAP1, the N-terminal domain of baculoviral IAP repeats (BIR), particularly the BIR1-2 domain (Fig. 7F-H and Fig. S11B-C). These results imply that LKB1 and JAK1 may compete for interacting with cIAP1. In support of this notion, the JAK1 level was decreased in the cIAP1 complex upon LKB1 overexpression (Fig. 7I). Altogether, these results confirm the LKB1-cIAP1 and cIAP1-JAK1 PPI and suggest that they are mutually exclusive.

To probe how the LKB1 status may impact its interaction with IAP, we examined the structural basis of LKB1 for its interaction with cIAP1. We found that the LKB1-cIAP1 interaction was primarily mediated by the LKB1 kinase domain, particularly the C-terminal lobe (132-347 aa) (Fig. 7J). In support of this notion, the LKB1-cIAP1 PPI was significantly compromised with LKB1 K78I, a kinase dead mutant (Fig. 7K). The naturally occurring LKB1 truncation mutants that lack the kinase domain were unable to interact with cIAP1 (Fig. 7L and Fig. S11D). These results suggest that the LKB1-cIAP1 PPI is LKB1 kinase-dependent, while cIAP1 may be dissociated from LKB1 in lung cancer cells with LKB1-inactivating mutations.

Due to the competitive nature of LKB1 and JAK1 for cIAP1 interaction, the LKB1 interaction state with cIAP1 may affect how cIAP1 regulates JAK1 and the IAP-dependency of cells. To test this contention, we utilized both genetic and pharmacological perturbation approaches to test the effect of the LKB1 and IAP1 status on JAK1 and its signaling. First, genetic knockdown of LKB1 led to a significant decrease in JAK1 protein, but not mRNA, suggesting a loss of JAK1 protein stability and the JAK1 function (Fig. 7M-N). In support of this notion, knockdown of LKB1 also impaired IFN*γ*signaling response as shown with reduced STAT1 phosphorylation (Fig. 7M), while overexpression of LKB1-WT sensitized IFN*γ*signaling response as measured by the STAT transcriptional reporter activity (Fig. 7O). Second, we found that birinapant, which degrades cIAP1 and thus disrupts cIAP1-JAK1 PPI, rescued the expression of JAK1 at protein, but not mRNA level, in LKB1-mut cells (Fig. 7P-Q and Fig. S11E-F). This birinapant-induced increase of JAK1 protein level is likely due to the stabilization of JAK1 protein through inhibition of cIAP1-associated ubiquitination (Fig. 7R and Fig. S11G). Further, upregulated JAK1 with the birinapant treatment was correlated with significant JAK1 downstream signaling activation, as shown by increased IFN*γ*-induced STAT1/3 phosphorylation (Fig. 7S and Fig. S11H), STAT transcriptional reporter activity (Fig. S11I-J), and the expression of target genes IRF1 and TAP1n (Fig. 7T-U and Fig. S11K-L). These results suggest that lung cancer cells with LKB1 mutations have rewired cIAP1-JAK1 PPI that leads to impaired IFN*γ*-JAK-STAT signaling.

Altogether, the identification of the LKB1 status controlled cIAP1-JAK1 regulatory axis provides molecular connectivity basis underpinning IAP- and STING-dependency in LKB1-mut lung cancer cells. Our data suggest that in LKB1-WT cells, LKB1 sequesters IAP1 from the JAK1 complex, maintaining a functional IFN*γ*-JAK-STAT signaling and the basal level of STING, while loss of LKB1 in LKB1-mut cells permits IAP1 to bind and degrade JAK1, impairing the JAK-STAT signaling and the STING downregulation. This IAP-induced JAK-STING pathway silencing can be pharmacologically reversed by small molecule IAP inhibitors (Fig. 7V).

## Discussion

Rewiring of tumor intrinsic circuitry enables cancer cell avoidance from immune attacks, ultimately leading to therapeutic failure (Kalbasi and Ribas, 2020; Litchfield et al., 2021; Wellenstein and de Visser, 2018). For example, lung adenocarcinomas harboring mutations in *LKB1* are highly aggressive, treatment refractory, and insensitive to immune checkpoint inhibitors (Calles et al., 2015; El Osta et al., 2019; Skoulidis et al., 2018; Xiao et al., 2016), representing a major clinical challenge. It is urgent to develop targeting strategies to enhance tumor response to therapies. Our studies with the chemical biology and molecular interaction approaches uncovered a mutated LKB1-created dependency of lung cancer cells on IAP and IAP-mediated downregulation of the JAK1-STING innate immunity pathway for the observed therapeutic resistance. Mechanistic examination revealed an intricate dynamic protein interaction hub by which LKB1 interacts with IAP in competition with JAK1, which offers a working model that the loss of LKB1 in LKB1-mut cancer cells allows IAP to engage JAK1 to control JAK1-mediated IFN*γ* response and the effector STING pathway, underpinning the IAP dependency (Fig. 7). In support of this model, pharmacological inhibition of IAP with birinapant synergizes with IFN*γ* to reverse STING expression and function to promote tumor cytotoxic T cell infiltration and induce apoptotic death of LKB1-mut lung cancer cells. Importantly, such an immune-dependent mode of action of IAP inhibitors in LKB1-mut cancer cells revealed by the HTiP approach is strongly supported by *in vivo* data. Birinapant effectively induced shrinkage of LKB1-mutant tumors in an immune-competent mouse model while showing no effect in an immune-deficient nude mouse model, presenting a potential therapeutic strategy for lung cancer patients with mutated LKB1.

The regulatory complex uncovered in our study couples the status of IAP to the activation state of the JAK1-STING pathway. Downregulation of tumor intrinsic STING expression has been frequently observed in many tumor types (Della Corte et al., 2020; Kitajima et al., 2019; Konno et al., 2018; Song et al., 2017; Xia et al., 2016a; Xia et al., 2016b). However, the regulatory mechanisms underlying STING expression control are context dependent and remain to be elucidated (Kitajima et al., 2019; Olagnier et al., 2018; Tsuchida et al., 2010; Wang et al., 2014). In the context of LKB1-mut LUAD, STING expression downregulation has been associated with transcriptional repression (Della Corte et al., 2020; Kitajima et al., 2019). Distinct from previously identified mechanism of epigenetic reprogramming mediated by DNMT1 and EZH2 (Kitajima et al., 2019; Morel et al., 2021), our data reveal a transcriptional program by which the IFN*γ*-JAK-STAT immune response signaling pathway dictates the STING expression in LKB1-mut cells. Our data further suggest that tumor intrinsic STING expression could be a potential biomarker for immune responsiveness, while restoring STING expression could be effective therapeutic strategies in LKB1-mut LUAD. Our study offers alternative approaches with the JAK1-directed mechanisms to reactivate the STING pathway in various tumors, which may be synergistic with the reported epigenetic modulation approach.

Dysregulation of IAP and JAK has been identified in a number of tumor types. For example, overexpression of IAPs has been linked to tumor progression, evasion of apoptosis and poor prognosis (Krajewska et al., 2003; Mizutani et al., 2007; Tamm et al., 2000). On the other hand, the loss-of-function JAK mutations have been associated with immune evasion and primary resistance to immunotherapy (Zaretsky et al., 2016). In contrast to the aberrant IAP and JAK function due to alterations at the genetic level, our data suggest that cIAP1 and JAK1 may be dysregulated at the protein level via rewired protein-protein interactions. Our study suggests that sequestration of IAP1 by LKB1 helps maintain a functional IFN*γ*-JAK-mediated immune response in LKB1-WT cells, whereas in LKB1-mut cells, IAP1 directly impinges on JAK1 and impairs tumor immune response. Thus, IAP’s pro-oncogenic activity could be enhanced by the loss of its bound LKB1 in LKB1 mutant cancer, while JAK1 could be directly weakened by IAP interaction, leading to STING suppression and immune resistance. These results implicate that such immune response regulatory mechanism at JAK1 protein level might be a general phenomenon in other tumor types with WT JAK1 (Patel et al., 2017). Moreover, the rewired JAK1 PPIs, such as IAP1-JAK1 in LKB1-mut cells, might be a promising therapeutic target, inhibition of which may sensitize tumors for enhanced immune response. Like the demonstrated IAP inhibitors, the IAP/JAK1 PPI antagonists may have the potential to re-activate the STING activity for therapeutic development.

The discovered IAP-JAK1 interaction suggests an emerging pathway for the action of IAP inhibitors as anticancer immune enhancers. IAP inhibitors, also known as SMAC mimetics, have been studied extensively as potential anti-tumor agents (Dineen et al., 2010; Krepler et al., 2013; Vince et al., 2007). IAP inhibitors were designed to block the inhibitory effect of IAP on caspase-mediated apoptosis signaling and thus promote cell death. Our data suggest that IAP inhibitors alone have minimal cell death-inducing effect on LKB1-mut tumors. However, immune factors, such as IFN*γ*, is required for IAP inhibitors to achieve their potent anti-tumor effect. IAP inhibitors synergize with IFN*γ*to activate RIPK1-dependent cell death pathway in a colon cancer cell model. In LKB1-mut LUAD tumors, however, the IAP inhibitor and IFN*γ* combination is JAK1-dependent involving the STING activation. Thus, the effect of the IAP inhibitor and IFN*γ* combination on its effector signaling could be context dependent, adding another layer of complexity to the emerging immunomodulatory activities of IAP inhibitors (Beug et al., 2017; Chesi et al., 2016; Clancy-Thompson et al., 2018; Dougan et al., 2010; Dougan and Dougan, 2018; Mo et al., 2019; Roehle et al., 2021).

Our results may have significant implications for the treatment of immune cold tumors. LKB1-mut LUAD has been characterized to exhibit an immune suppressive phenotype with multiple “immune cold” features, such as low PD-L1 expression, low tumor mutational burden, altered proinflammatory cytokine profiles, reprogrammed immune infiltration, remodeled extracellular matrix, and recently demonstrated STING expression downregulation (Cristescu et al., 2018; Della Corte et al., 2020; Gao et al., 2010; Gilbert-Ross et al., 2017; Kadara et al., 2017; Kitajima et al., 2019; Koyama et al., 2016; Skoulidis et al., 2015; Skoulidis et al., 2018). Therefore, effective strategies that re-inflame LKB1-mut tumors may have significant therapeutic potential. Indeed, the IAP inhibitors described here can restore STING expression, reinvigorate STING-mediated DNA-sensing pathway, re-induce innate immunity cytokine and chemokine production, promote chemotaxis of immune cells, and re-sensitize tumors for enhanced immune responsiveness. Our data suggest that IAP inhibitors may be promising immunotherapy adjuvants with immune checkpoint inhibitors or STING agonists to overcome LKB1-mut associated immune resistance.

## Significance

To advance precision immunotherapy, our study identifies IAP inhibitors as potent immune-dependent anti-tumor agents against LKB1-mut lung cancer cells, revealing an intrinsic IAP-dependency mechanism for immune escape. The inhibition of IAP promotes tumor cytotoxic immune cell infiltration and induces drastic shrinkage of LKB1-mutant tumors selectively in an immune-competent mouse model, demonstrating its therapeutic potential. The mode-of-action studies uncover a hidden IAP-JAK-STING regulatory axis that underpins the IAP dependency. The discovery of LKB1-cIAP1-JAK1 trimolecular complexes provides a molecular basis that connects the LKB1-mut genotype with its immune-suppression phenotype. These data support the clinical investigation of using IAP inhibitors as immunotherapy adjuvants and tumor-intrinsic STING expression as a biomarker to augment the LKB1-mut tumor responsiveness to immunotherapies, such as checkpoint inhibitors or STING-targeted agents, to accelerate oncogenic mutation-driven precision medicine.

## Acknowledgements

This work was supported by the National Cancer Institute’s Office of Cancer Genomics Cancer Target Discovery and Development (CTD^2^) initiative network (U01CA217875 to HF), the NCI Emory Lung Cancer SPORE (P50CA217691 to SR, HF) Career Enhancement Program (XM, AAI; P50CA217691), the NCI R01 (R01CA203928 to WZ), the NCI R37 (R37CA255459 to XM), the Imagine, Innovate and Impact (I^3^) Funds from the Emory School of Medicine and through the Georgia CTSA NIH award (UL1-TR002378), the Winship Invest$ Team Science award, and Winship Cancer Institute (NIH 5P30CA138292). CS is a visiting student in the Emory University School of Medicine-Xiangya Medical School student exchange program. AW is a visiting student in the Emory University School of Medicine-Xi’an Jiaotong University Health Science Center student exchange program. We thank all members of the Fu lab for technical support and comments. We thank Rui Liu at Department of Pediatrics, Emory University School of Medicine, for her help with RNAseq data analysis. We acknowledge Emory University Integrated Cellular Imaging Core Shared Resources.

## Author Contributions

Conceptualization, X.M., and H.F.; *In vitro* molecular and cellular biology studies, C.S., X.M., Q.N., D.C., S.D., D.F., X.Z. and Y.D.; *In vivo* animal studies, R.J., X.M., C.S., W.Z.; Immunohistochemistry studies, R.J., W.Z., J.C., T.O., S.R., and G.S.; Mass cytometry studies, D.B.D., K.M.D. and M.V.D.; Informatics analysis, A.W. and A.A.I.; Data Analysis, C.S., X.M., R.J. W.Z., K.M.D., J.C., T.O., S.R., G.S., and H.F.; C.S., X.M., and H.F. wrote the initial manuscript; Funding acquisition, H.F.; Resources, W.Z. and H.F.; Supervision, W.Z., X.M. and H.F. All were involved in editing.

---

**Figure S1.**
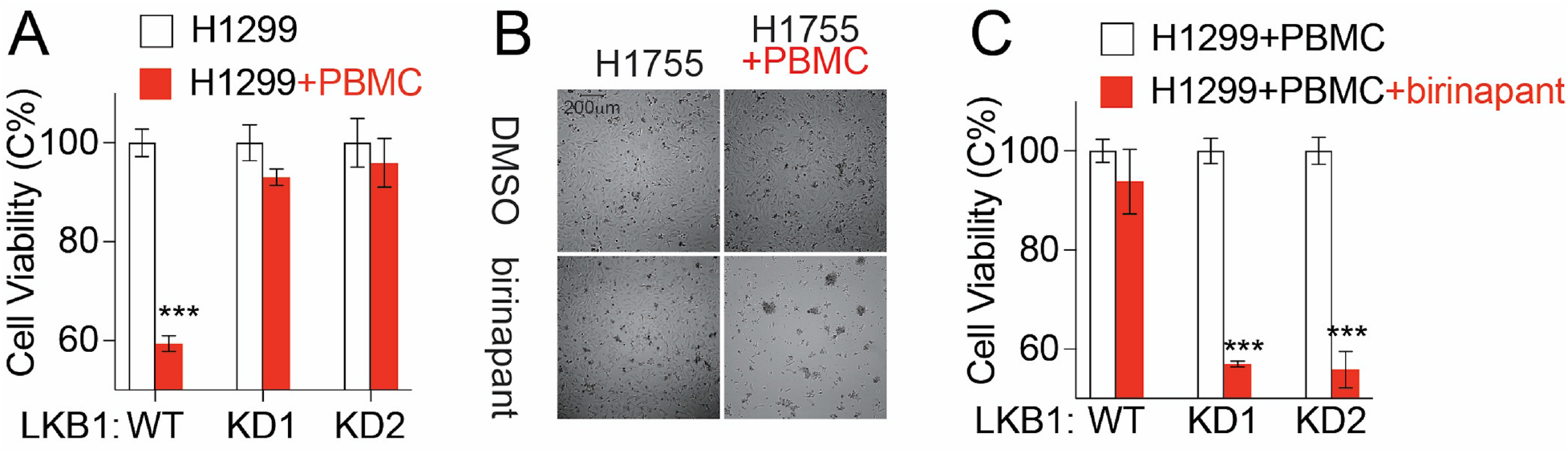
Birinapant-induced sensitization of immune responsiveness in LKB1-mut cells, related to Fig. 1. **(A)** Bar graph showing immune resistence in LKB1-knockdown (KD) H1299 cells. Parental H1299 (WT) or isogenic LKB1-knockdown (KD1 and KD2) cells were cultured alone or co-cultured with PBMC (E:T=15:1). The data are presented as mean±SD from three independent experiments. ***p*≤*0.001. **(B)** Representative images showing birinapant-induced sensitization of immune responsiveness in H1755 cells. H1755 cells were co-cultured with PBMC (E:T=1:1) for 4 days with or without birinapant(500nM) treatment as indicated. **(C)** Bar graph showing birinapant-induced sensitization of immune responsiveness in LKB1-mut cells. Parental H1299 (WT) or isogenic LKB1-knockdown (KD1 and KD2) cells were co-cultured with PBMC (E:T=15:1) and treated with birinapant (150 nM) for 4 days. The data are presented as mean±SD from three independent experiments. ***p*≤*0.001.

**Figure S2.**
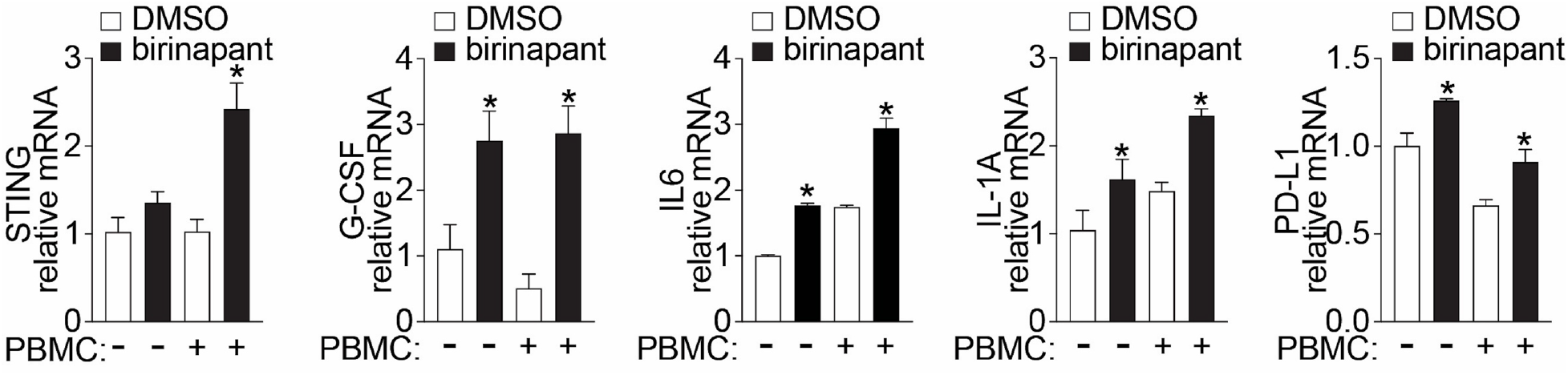
Birinapant-induced immune-dependent STING expression in LKB1-mut cells, related to Fig. 2. Bar graph showing STING, G-CSF, IL6, IL-1A and PD-L1 relative mRNA expression in H1755 cancer cells that were cultured alone or co-culture with immune cells (E:T=1:1). The change of relative mRNA expression of the selected genes was expressed as fold-of-change upon birinapant (50 nM) treatment over DMSO control. The data are presented as mean±SD from three independent experiments. *p*≤*0.05.

**Figure S3.**
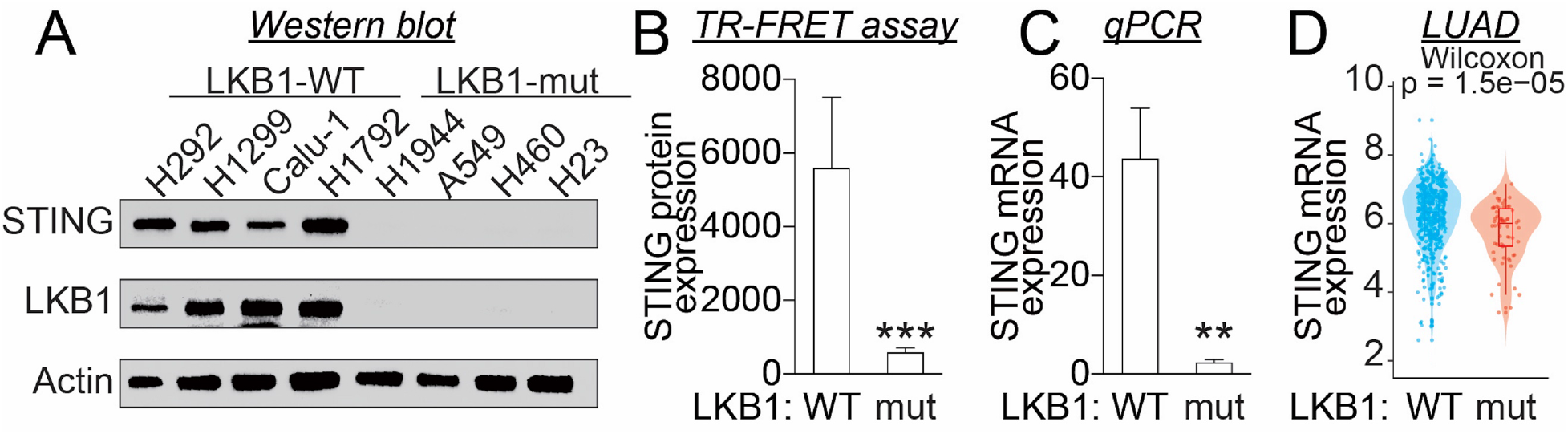
Downregulation of STING expression in LKB1-mut cells, related to Fig. 2. **(A)** Western blot and **(B)** TR-FRET assay analysis of STING protein expression in LUAD cell lines with LKB1-WT (Calu-1, H1299, H1792 and H292) or LKB1-mut (A549, H1944, H23 and H460). **(C)** qPCR assay and **(D)** TCGA analysis of STING mRNA level in LUAD cell lines and patient samples. WT: wild-type LKB1; mut: inactivating mutations of LKB1. The data in (B-C) are presented as mean±SD of multiple cell lines. **p*≤*0.01, ***p*≤*0.001.

**Figure S4.**
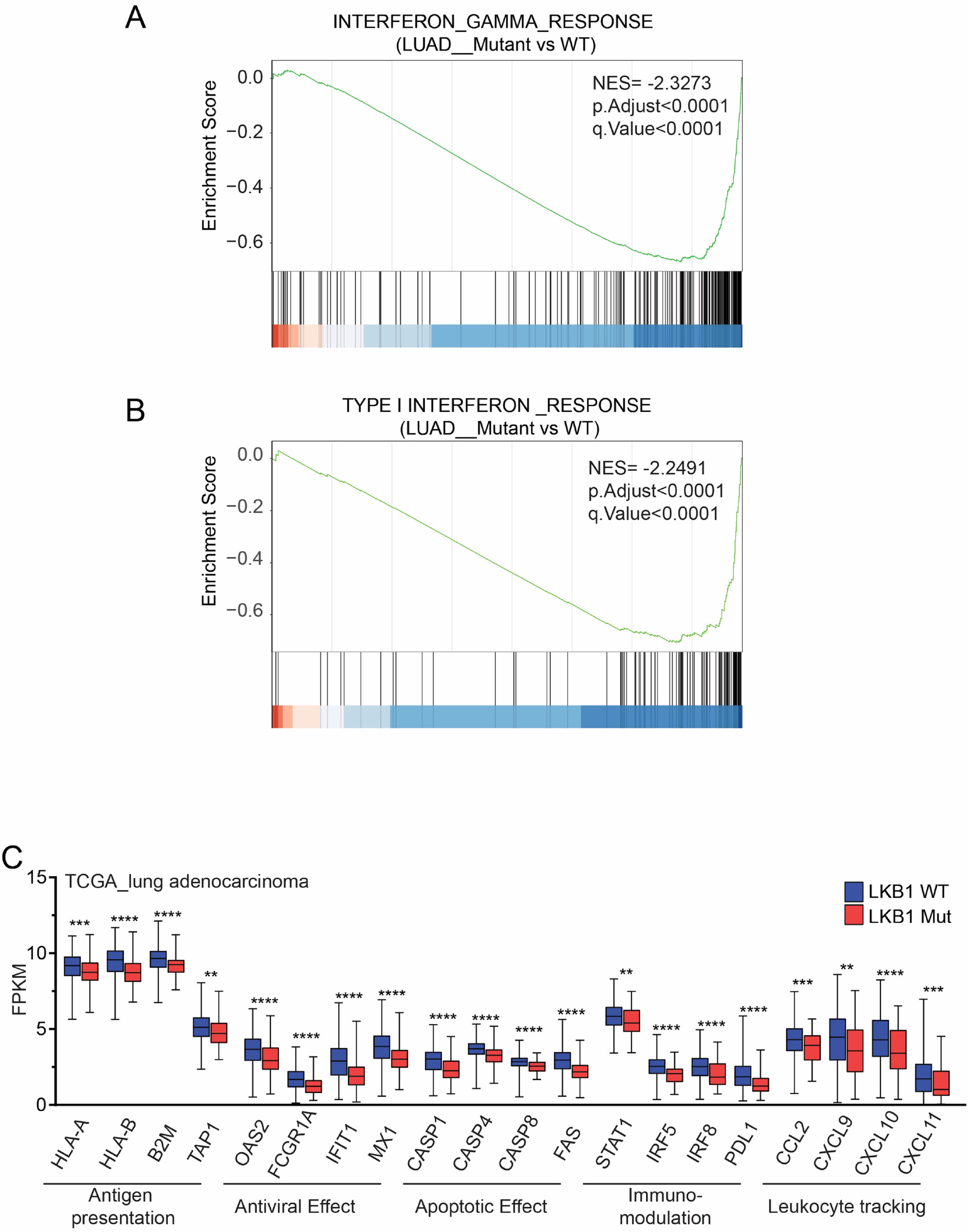
Downregulation of type I interferon and IFNγ pathway in LKB1-mut cells, related to Fig. 3. **(A-B)** Gene set enrichment analysis showing significantly enrichment of differential expression genes from LKB1-mut as compared to LKB1-WT LUAD TCGA samples. **(C)** Downregulation of type I interferon and IFN*γ* pathway-associated gene in LKB1-mut LUAD TCGA samples. **p*≤*0.01, ***p*≤*0.001, **** p*≤*0.0001.

**Figure S5.**
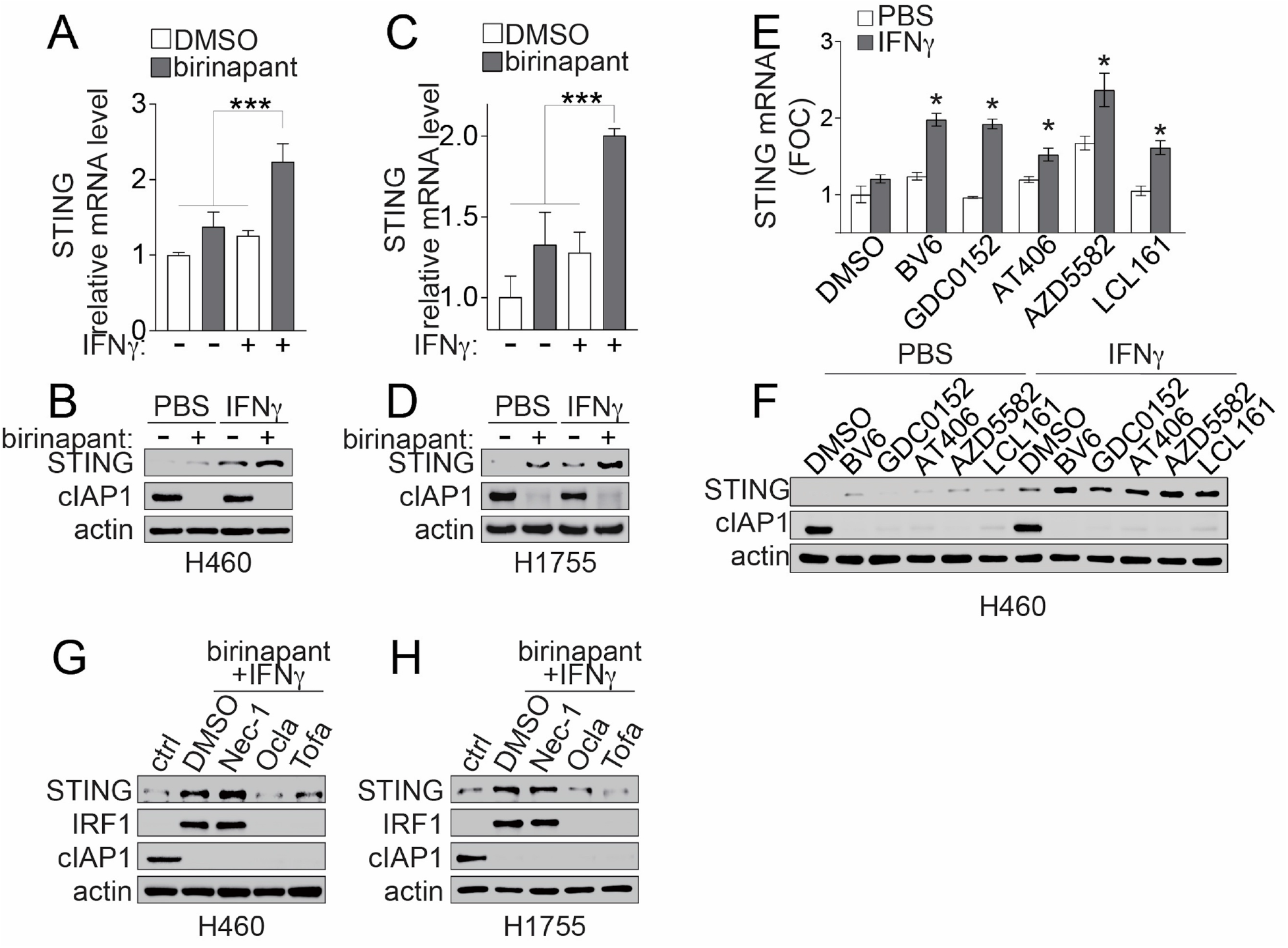
IAP inhibitor-induced IFNγ-dependent STING expression in LKB1-mut cells, related to Fig. 3. **(A-F)** STING mRNA (A, C and E) and protein (B, D and F) expression in LKB1-mut cells upon treatment of IAP inhibitor, IFN*γ*, or in combination. H460 (A-B) and H1755 (C-D) cells were treated with birinapant (500 nM) or other IAP inhibitors (500 nM), IFN*γ* (5 ng/mL), or in combination as indicated for 24 hours. STING relative mRNA expression was expressed as fold-of-change over untreated DMSO control and presented as mean±SD of three independent experiments. STING protein expression was analyzed by SDS-PAGE and western blot with indicated antibodies and was presented as representative immunoblots. *p*≤*0.05, ***p*≤*0.001. **(G-H)** Representative immunoblot showing STING protein expression in LKB1-mut cells upon birinapant and IFN*γ*combination treatment with additional JAK inhibitors. H460 (G) and H1755 (H) cells were treated with birinapant (500 nM) plus IFNg (5 ng/mL) in combination with JAK inhibitors, oclacitinib (Ocla, 10 μM) and tofacitinib (Tofa, 10 μM), or RIPK inhibitor, necrostatin-1 (Nec-1, 10 μM), as indicated. STING protein expression was analyzed by SDS-PAGE and western blot with indicated antibodies.

**Figure S6.**
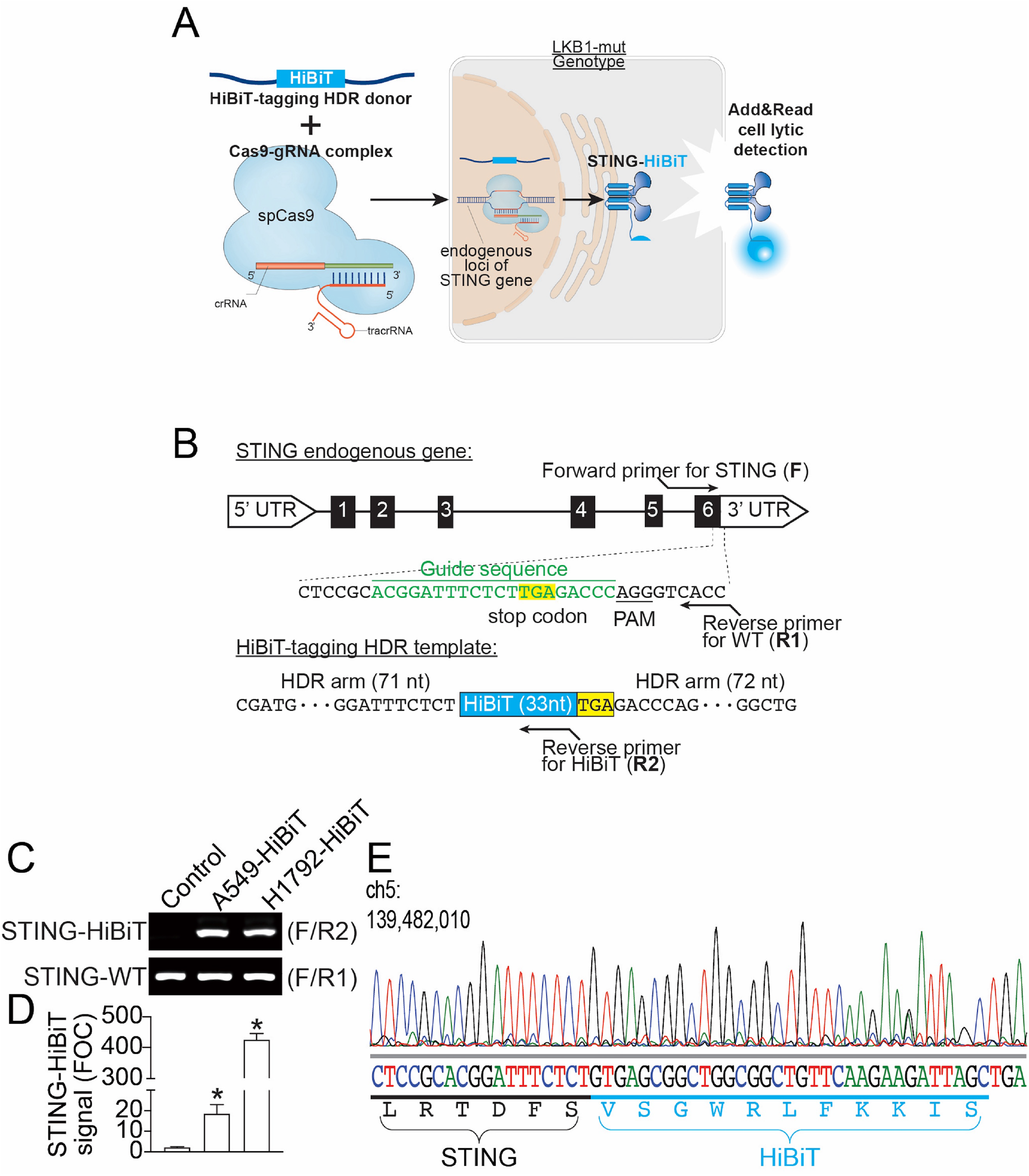
Design and development of STING-HiBiT assay to monitor endogenous STING expression, related to Fig. 3. **(A)** Schematic illustration of CRISPR/Cas9-HiBiT tagging technology for STING-HiBiT cell engineering. **(B)** Schematic illustration of gRNA and HDR template design. **(C-E)** Validation of STING-HiBiT engineered cells by PCR using primer pair as indicated (C), HiBiT luminescence (D) and sanger sequencing (E). The data in (D) are presented as mean±SD of three independent experiments. *p<0.05.

**Figure S7.**
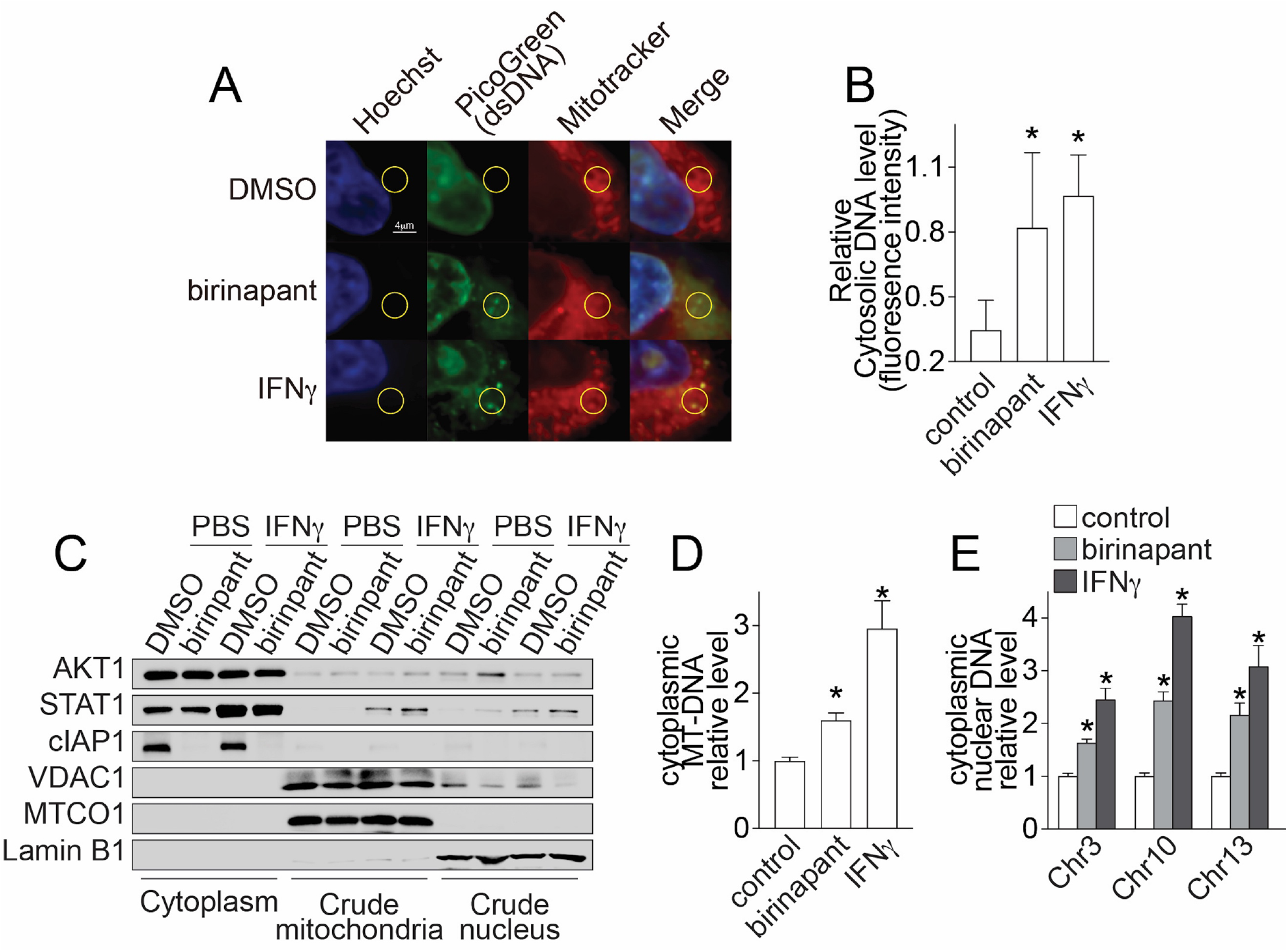
Birinapant- and IFNγ-induced increase of cytosolic DNA in LKB1-mut cells, related to Fig. 4. **(A)** Representative fluoresence images showing cytosolic DNA staining in A549 cells. A549 cells were treated with birinapant (500 nM) or IFN*γ* (1 ng/mL) for 24 hours and immunostained with antibodies as indicated. **(B)** Bar graph showing cytosolic DNA quantification from immunostaining. Relative fluorescence intensity was caculated as the ratio of PicoGreen intensity to Mitotracker intersity. The data are presented as as mean±SD of the fluoresence intensity from thirty area of interest. *p<0.05. **(C)** Representative immunoblot showing the successful preparation of lysate from cytoplasm. A549 cells were treated with birinapant (500 nM) or IFN*γ* (1 ng/mL) for 24 hours and subjected to cell lysate preparation from cytoplasm, mitochondria, and nucleus. Cell lysate was confirmed using representative subcellular compartment resident proteins, such as AKT1 in cytoplasm, VDAC1 and MTCO1 in mitochondria, and Lamin B1 in nucleus. **(D-E)** Bar graphs showing birinapant- and IFN*γ*-induced increase of cytoplasmic DNA released from mitochondria (D, MT-DNA) or nucleus (E, nuclear DNA). Cytoplasmic DNA was quantified by qPCR using MT-DNA or nuclear-DNA specific primers.

**Figure S8.**
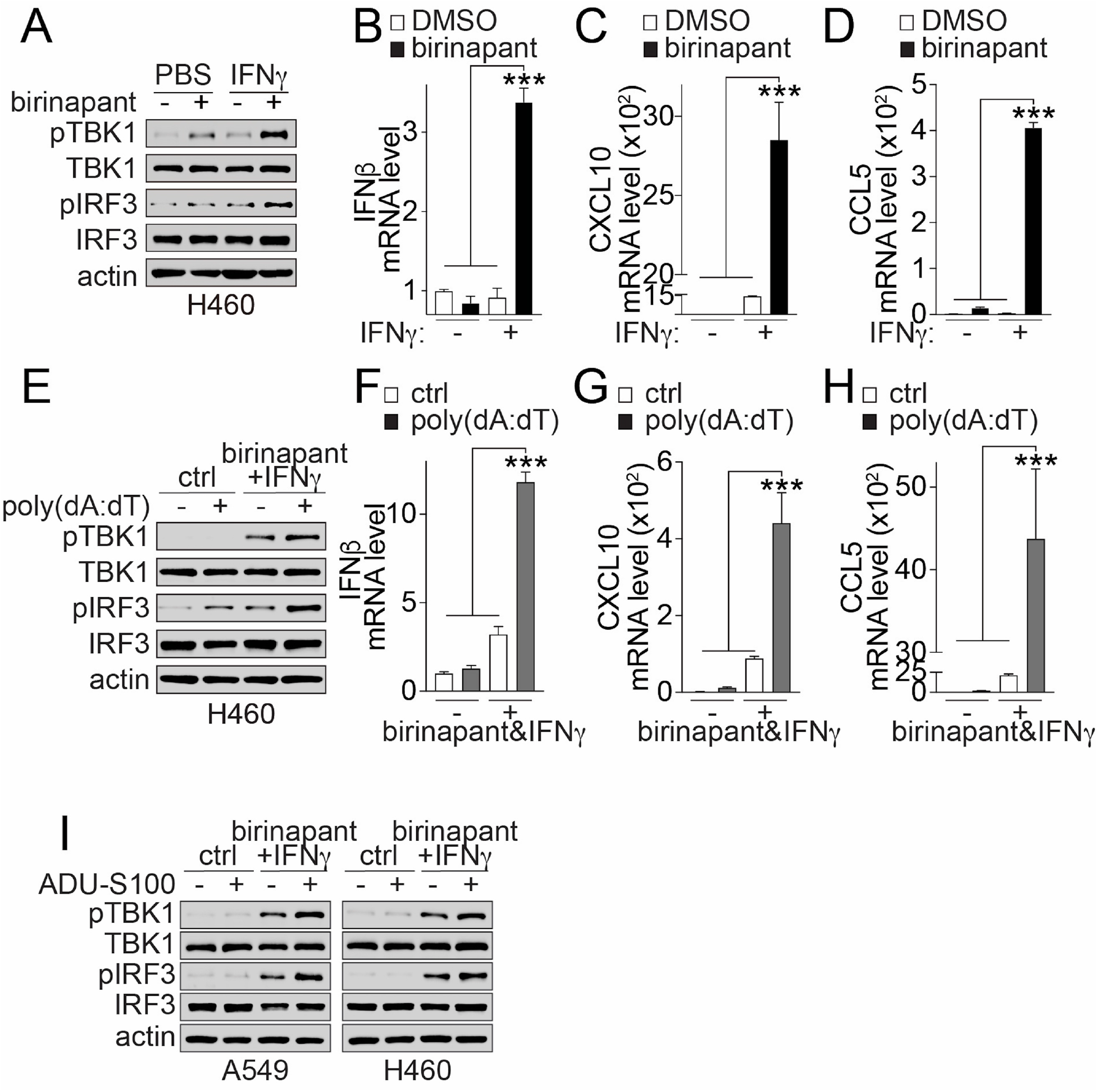
Birinapant synergizes with IFNγ to induce STING-mediated DNA sensing pathway activation in LKB1-mut cells, related to Fig. 4. **(A)** Representative immunoblot showing birinapant and IFN*γ*combination-induced activation of TBK1 and IRF3. H460 cells were treated with birinapant (500 nM) and/or IFN*γ*(5 ng/mL) for 24 hours as indicated. Cell lysate were analyzed by SDS-PAGE and western blot with antibodies as indicated. **(B-D)** Bar graphs showing birinapant and IFN*γ* combination-induced expression of IFN*β* (B), CXCL10 (C) and CCL5 (D). H460 cells were treated with birinapant (500 nM) and/or IFN*γ* (5 ng/mL) for 24 hours as indicated. Gene expression were analyzed by qPCR and presented as mean±SD from three independent experiments. *** p*≤*0.001. **(E)** Immunoblot showing poly(dA:dT)-induced TBK1 and IRF3 activation. H460 cells were treated with poly(dA:dT) (1 μg/mL) for 4 hours in the presence or absence of 24-hour pre-treatment with birinapant (500 nM) and IFN*γ* (5 ng/mL) combination as indicated. Cell lysate were analyzed by SDS-PAGE and western blot with antibodies as indicated. **(F-H)** Bar graphs showing poly(dA:dT)-induced expression of IFN*β* (F), CXCL10 (G) and CCL5 (H). H460 cells were treated with poly(dA:dT) (1 μg/mL) for 4 hours in the presence or absence of 24-hour pre-treatment with birinapant (500 nM) and IFN*γ*(5 ng/mL) combination as indicated. Gene expression were analyzed by qPCR and presented as mean±SD from three independent experiments. *** p*≤*0.001. **(I)** Representative immunoblot showing ADU-S100-induced activation of TBK1 and IRF3 phosphorylation. A549 or H460 cells were treated with ADU-S100 (10 μM) for 3 hours in the presence of 24-hour pre-treatment of birinapant (500 nM) plus IFN*γ*(1 and 5 ng/mL for A549 and H460 cells, respectively). Cell lysate were analyzed by SDS-PAGE and western blot with antibodies as indicated.

**Figure S9.**
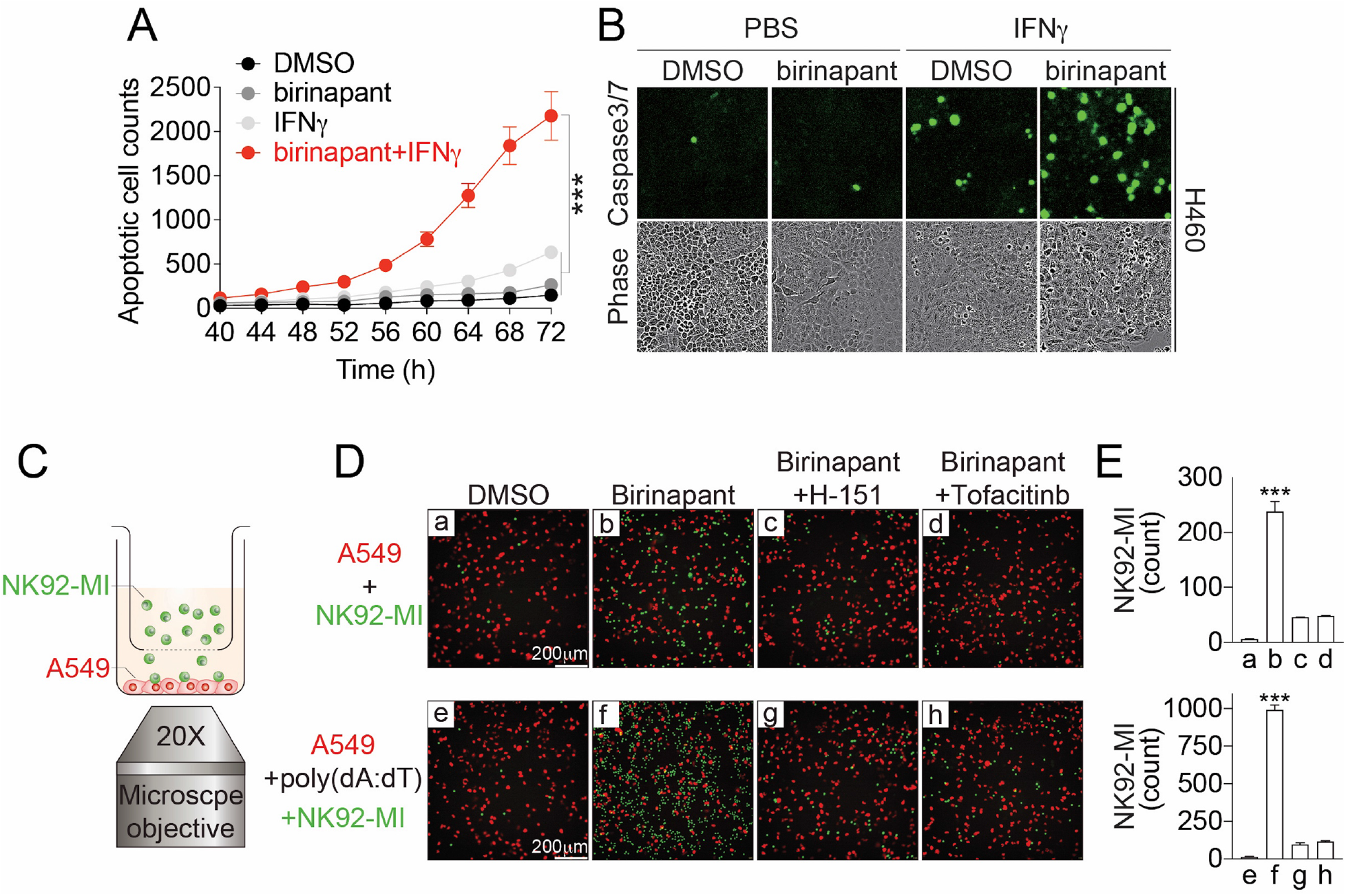
Birinapant induces STING-mediated apoptosis of LKB1-mut cancer cells and migration of immune cells *in vitro*, related to Fig. 5. **(A)** Time-dependent curve of birinapant-induced cell apoptosis. H460 cells were treated with birinapant (500 nM), IFN*γ* (5 ng/mL) or in combination as indicated. Cell apoptosis was measured and monitored in real-time using DEVD-based fluorogenic caspase-3/7 apoptosis reporter assay up to 72 hours. The data were expressed as the count of green fluorescence positive cells and presented as mean±SD from three independent experiments. *** p*≤*0.001. **(B)** Representative images showing birinapant-induced cell apoptosis. H460 cells were treated with birinapant (500 nM), IFN*γ* (5 ng/mL) or in combination as indicated. Green fluorescence (caspase-3/7 apoptosis reporter) and phase-contrast images were acquired using IncuCyte. **(C)** Schematic illustration of transwell assays for measuring immune cell infiltration *in vitro*. **(D)** Representative fluorescence images showing birinapant-induced NK92-MI cells migration. A549 cells and NK92-MI cells were co-cultured in transwell as shown in (C) and were treated with birinapant (100 nM), poly(dA:dT) (1 μg/mL), or in combination with STING antagonist, H151 (5 μM), or JAK inhibitor (JAKi), tofacitinib (10 μM), for 48 hours as indicated. Fluorescence images of A549 cells (red) and infiltrated CD56^+^ NK92-MI cells (green) were acquired at the endpoint using ImageXpress Micro high-content imaging system. **(E)** Bar graph showing the quantification of infiltrated NK92-MI cells in transwell-based migration assays. The data were expressed as the count of green fluorescence positive NK92-MI cells and presented as mean±SD from three independent experiments. *** p*≤*0.001.

**Figure S10.**
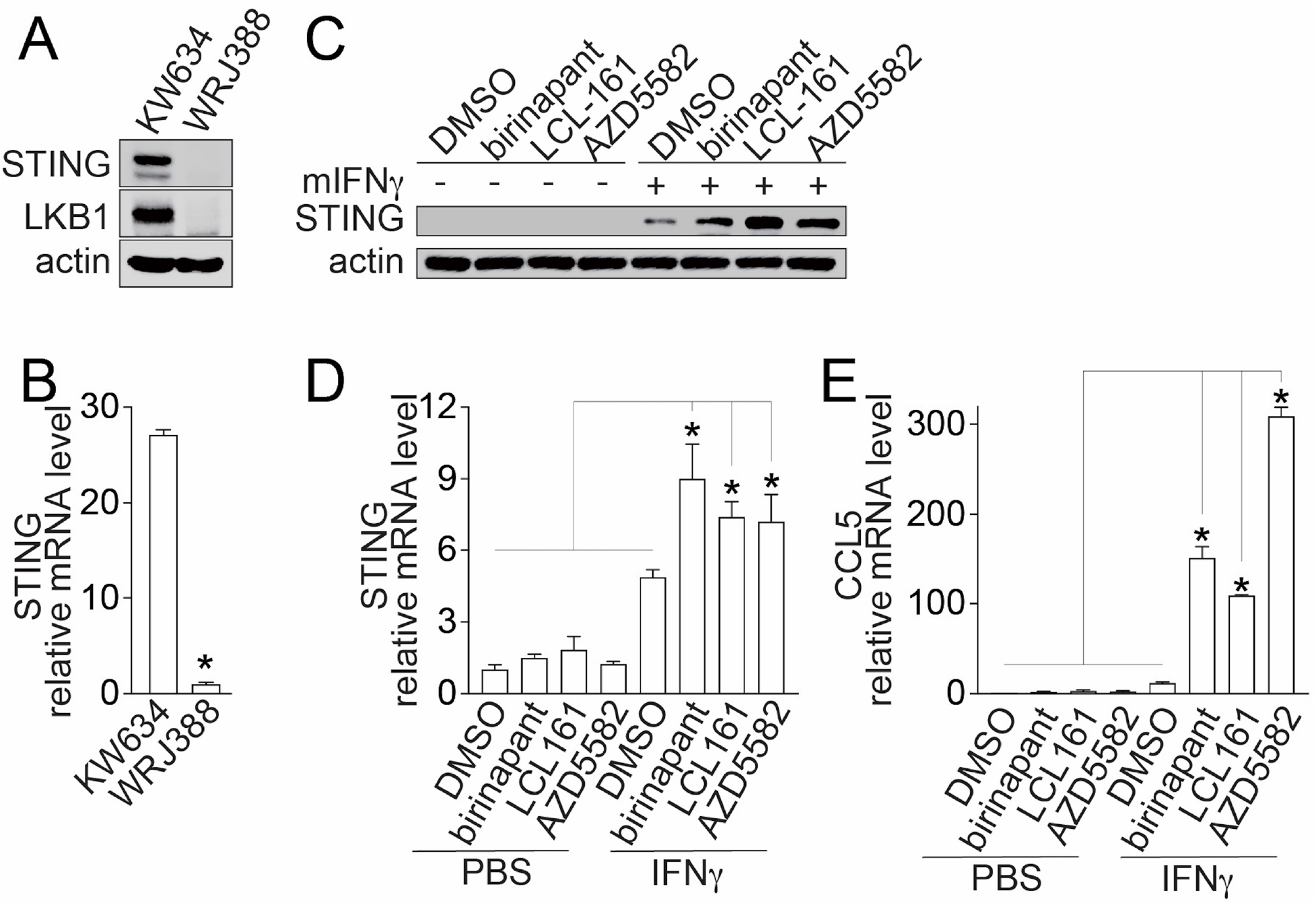
Birinapant synergize with mouse IFNγ to induce STING expression and signaling activation in *Lkb1*-mut mouse lung cancer cells *in vivo*, related to Fig. 6. **(A-B)** Representative immunoblot and bar graph showing STING protein (A) and mRNA (B) expression downregulation in WRJ388, a *Lkb1*-mut mouse lung cancer cells, as compared with KW634, a *Lkb1*-WT mouse lung cancer cells control. **(C-E)** Representative immunoblot and bar graph showing STING protein expression (C) and STING (D) or CCL5 (E) mRNA expression in WRJ388 cells treated with IAP inhibitors (500 nM) and mouse IFN*γ* (mIFN*γ*, 10 ng/mL) as indicated for 24 hours. STING protein expression was analyzed by SDA-PAGE and western blot with antibodies indicated. The relative STING and CCL5 mRNA expression was analyzed by qPCR, expressed as fold-of-change over untreated DMSO control and presented as mean±SD from three independent experiments. * p*≤*0.05.

**Figure S11.**
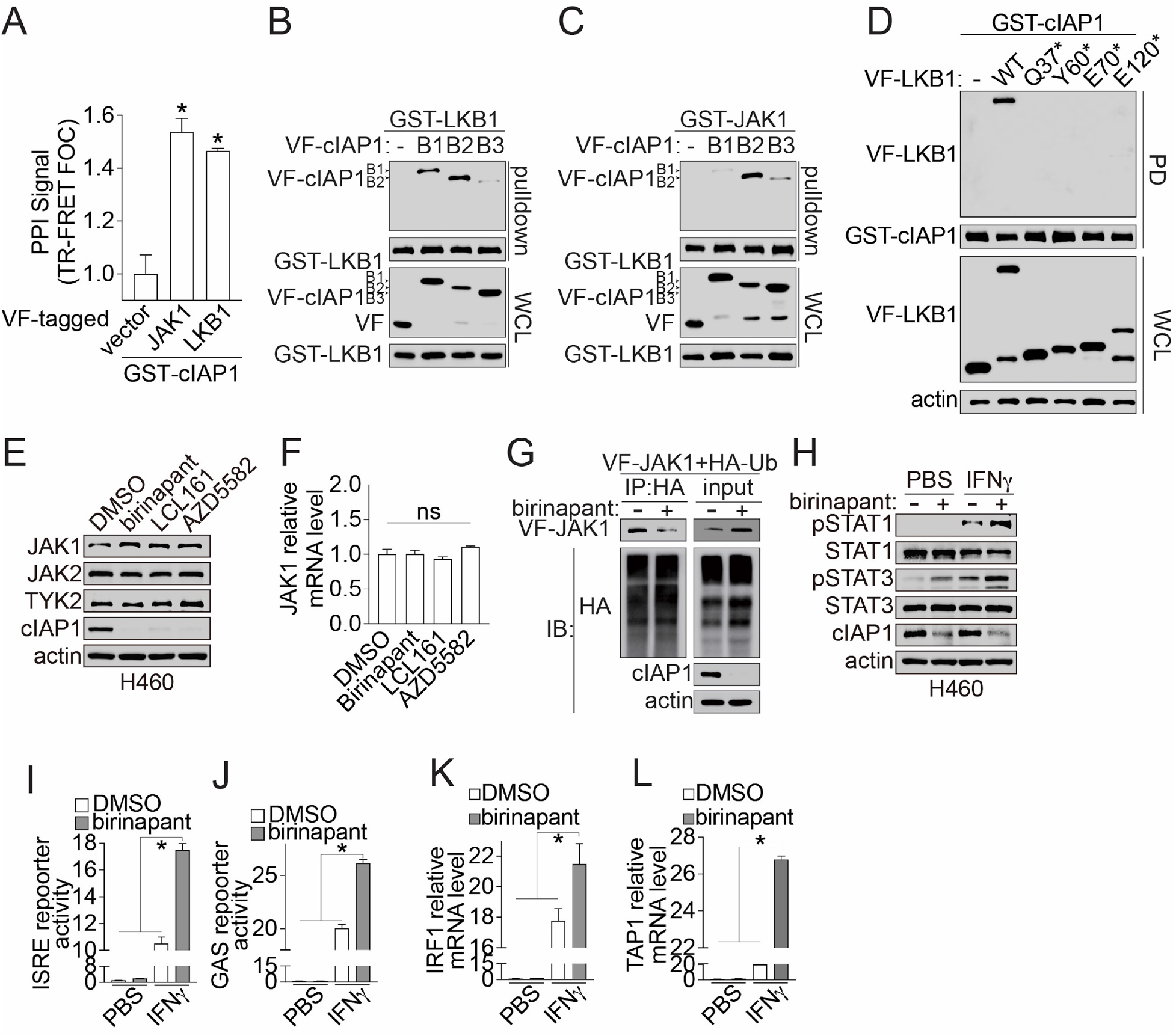
Identification and characterization of LKB1-cIAP1-JAK1 trimolecular interaction, related to Fig. 7. **(A)** Bar graph showing PPI signal between cIAP1 with LKB1 and JAK1. PPI signal were measured using cell lysate from HEK293T cells co-expressing GST-cIAP1 and Venus-flag (VF)-tagged LKB1 or JAK1 as indicated. The PPI signal were expressed as the fold-of-change over the corresponding empty vector controls and presented as the average of three technical replicates from the primary screen. **(B-C)** Immunoblot showing mapping of LKB1- (G) and JAK1- (H) binding domain on cIAP1. Cell lysate from HEK293T cells co-expressing GST-LKB1 or JAK1 with and Venus-flag (VF)-tagged cIAP1 BIR1 domain truncation (B1), BIR2 domain truncation (B2), and BIR3 domain truncation (B3) were subjected to the GST-pulldown as indicated. The pulldown complex and whole cell lysate (WCL) were analyzed by SDS-PAGE and western blot with antibodies as indicated. **(D)** Representative immunoblot showing PPI signal between cIAP1 and LKB1 WT or naturally occuring mutant. Cell lysate from HEK293T cells co-expressing GST-cIAP1 and Venus-flag (VF)-tagged LKB1 WT or naturally occuring mutant as indicated were subjected to the GST-pulldown assay. The pulldown complex and whole cell lysate (WCL) were analyzed by SDS-PAGE and western blot with antibodies as indicated. **(E-F)** JAK1 protein (E) and mRNA (F) expression upon IAP inhibitor treatment in LKB1- mut cells. H460 cells were treated with IAP inhibitors (500 nM) as indicated for 24 hours. JAK1 protein level was measured using cell lysates and were analyzed by SDS-PAGE with antibodies as indicated. The relative JAK1 mRNA expression was measured using qPCR. The data are expressed as fold-of-change over the DMSO control (FOC) and presented as mean±SD from three independent experiments. ^ns^p>0.05 and FOC<1.5. **(G)** Representative immunoblot showing birinapant-induced decrease of ubiquitinated JAK1. Cell lysate from HEK293T cells expressing Venus-flag(VF)-tagged JAK1 and HA-tagged ubiquitin (Ub) with (+) or without (-) 16-hour birinapant (500 nM) plus 6-hour MG132 (20 μM) treatment were subjected to HA-beads immunoprecipitation. Immunoprecipitated (IP) complex and whole cell lysate input were analyzed by SDS-PAGE and western blot with antibodies as indicated. **(H)** Immunoblot showing birinapant-induced sensitization of IFN*γ*-mediated STAT1/3 activation in LKB1-mut cells. H460 cells were treated with IFN*γ* (5 ng/mL) or PBS control for 20 minutes with (+) or without 24-hour pretreatment of birinapant (500 nM). Cell lysate were analyzed by SDS-PAGE and western blot with antibodies as indicated. **(I-J)** Bar graphs showing birinapant-induced increase of STAT-driven transcriptional luciferase reporter activity. A549 cells stably expressing ISRE-luc (I) or GAS-luc (J) reporter plasmid were treated with birinapant (500 nM), IFN*γ* (1 ng/mL) or in combination as indicated. The reporter signal was measured by firefly luciferase kit, expressed as fold-of-change over the untreated DMSO and PBS control, and presented as as mean±SD from three independent experiments. *p*≤*0.05. **(K-L)** Bar graphs showing birinapant-induced sensitization of IFN*γ*-mediated IRF1 (K) and TAP1 (L) mRNA expression. H460 cells were treated with birinapant (500 nM), IFN*γ* (5 ng/mL), or in combination as indicated for 24 hours. The relative IRF1 and TAP1 mRNA expression was measured using qPCR. The data are expressed as fold-of-change over the PBS control (FOC) and presented as mean±SD from three independent experiments. *p*≤*0.05.

## STAR METHODS

### RESOURCE AVAILABILITY

#### Lead Contact

Further information and requests for resources and reagents should be directed to and will be fulfilled by the Lead Contact, Haian Fu.

#### Materials Availability

Plasmids generated in this study are available upon request to the Lead Contact. Other materials are available through commercial sources (see Key Resource Table)

#### Data and Code Availability

Analyzed small molecule screening data sets and mRNA sequencing data sets are available through CTD^2^ data portal (https://ocg.cancer.gov/programs/ctd2/data-portal).

### EXPERIMENTAL MODEL AND SUBJECT DETAILS

All cell lines were incubated at 37°C in humidified conditions with 5% CO_2_. Human embryonic kidney 293T cells (HEK293T; ATCC, CRL-3216) were maintained in Dulbecco’s Modified Eagle’s Medium (DMEM; Corning, #10-013-CV). Human lung adenocarcinoma cell lines, including A549 (ATCC, CCL-185), Calu-1 (ATCC, HTB-54), H1299 (ATCC, CRL-1848), H1755 (ATCC, CRL-5892), H1792 (ATCC, CRL-5895), H1944 (ATCC, CRL-5907), H23 (ATCC, CRL-5800), H292 (ATCC, CRL-1848), and H460 (ATCC, HTB-177), were cultured in Roswell Park Memorial Institute(RPMI) 1640 medium. The isogenic LKB1-wildtype and mutant cells were generated by lentivirus-transduction of corresponding shRNA or cDNA plasmids in the parental cells. Mouse lung adenocarcinoma cells, including WRJ388 (Jin et al., 2021) and KW634 (generous gifts from Dr. Kwok-Kin Wong) (Gandhi et al., 2009), were cultured in RPMI-1640 medium. Human immune cells, including Jurkat T cell (ATCC, TIB-152), NK92-MI (ATCC, CRL-2408), peripheral blood mononuclear cells (normal PBMC, ATCC, PCS-800-011), CD8^+^ T cells (Stemcell, 200-0164) and CD56^+^ NK cells (Stemcell, 70037) were cultured in RPMI-1640 medium. Cell culture medium was supplemented with 10% fetal bovine serum (ATLANTA biologicals, #S11550) and 100 units/ml of penicillin/streptomycin (Cell Gro, Cat# 30-002-CI).

All animal studies were approved and conducted according to the Emory University Institutional Animal Care and Use Committee (IACUC) guidelines. Five- to 6-week-old male athymic nude mice (18–20 g) were purchased from Harlan Laboratories. Syngeneic immune-competent *KL* (*Kras*^G12D^/*Lkb1*^-/-^) male mice were generated as described previously (Gilbert-Ross et al., 2017; Jin et al., 2021).

### METHOD DETAILS

#### Plasmids

Plasmids for mammalian expression of Glutathione S-transferase- (GST), Venus- Flag- (VF) and Human influenza hemagglutinin- (HA) tagged proteins were generated using the Gateway cloning system (Invitrogen, Waltham, MA, USA) according to the manufacturer’s protocol. The WT STK11 (LKB1) gene in pDONR223 was purchased from OpenBiosystem Kinome Entry ORF set. Other genes in pDONR221 were gifted from Drs. Gordon Mill and Yiu Huen Tsang at Oregon Health Science University. HA-Ubiquitin was purchased from Addgene (#18712) (Kamitani et al., 1997). The LKB1 and cIAP1 domain truncations in pDONR223 were generated using PCR and Gateway cloning system. The LKB1 mutations, including K78I and nonsense mutations, were introduced using QuikChange Lightning Site-Directed Mutagenesis Kit (Agilent Technologies, Cat# 210518). STING and LKB1 cDNA were cloned into pHAGE lentiviral vector for lentivirus packaging. Plasmids for shRNA knockdown were purchased from MISSON shRNA library (Sigma-Aldrich, see Key resource table). All the plasmids were confirmed by sequencing.

#### Transfection

Polyethylenimine(PEI, Cat#23966) transfection reagent was used for plasmid transfection in HEK293T cell. FuGene HD (Roche, Cat# E2920) was used in a ratio of 3 μl to 1 μg DNA for transfection in other cancer cells.

#### Immune and tumor cell co-culture assay

PBMCs, primary CD8^+^ T cells or CD56^+^ NK cells as effector immune cells and various human lung adenocarcinoma cells as target cells were used for co-culture assays. Tumor cells were seeded at specific density in 384-well cell culture plate (Corning #3764). Twenty-four hours later, effector were then thawed and co-cultured in RPMI-1640 medium with tumor cells in a dose dependent manner for four days. CD3 monoclonal antibody (100 ng/mL, OKT3, ThermoFisher) and human recombinant interleukin-2 (10 ng/mL, PeproTech) were used as activation cocktail to activate immune cells.

#### Cell proliferation measurement

The co-culture assay plates were imaged using the IncuCyte® S3 Live-Cell Analysis System (Essen Biosciences). The cancer cell proliferation was monitored and characterized as the percentage of confluence using the IncuCyte® basic analysis module. Because of the size distinction between effector immune cells and target cancer cells, the area filter of >400 μm^2^ was used to select cancer cells that are larger in size.

#### Cell Viability measurement

Cell Titer Blue (Promega, G8081) was added to each well. The plates were incubated for desired time at 37 °C to allow the generation of sufficient signal within the linear range. The fluorescence intensity of each well was read using an PHERAstar FSX multi-mode plate reader (Ex 545 nm, Em 615 nm; BMG LABTECH). Cells containing medium or immune cells alone were used as blank control for background correction.

#### High-throughput immunomodulator phenotypic (HTiP) screen

The primary HTiP screen was performed as described previously (Mo et al., 2019). Briefly, parental H1755 cells harboring LKB1 mutation were seeded in 384-well cell culture plate (2,000 cells/well in 40 μl medium; Corning, Cat#3764) and co-cultured with PBMCs (1,000 cells/well in 10 μl media containing the activation cocktail). The 2,036 Emory Enriched Library (EEL) compounds (100 nl) were used (Du et al., 2020; Mo et al., 2019; Tang et al., 2021). Compunds were added into wells in each plate using Biomek NXP Automated Workstation (Beckman) from a compound stock plate to give the final concentration of 2 μM. A parallel screening was performed with H1755 cells alone (2000 cells/well) in 50 μl medium containing the same amount of activation cocktail in the absence of PBMCs. After 4 days of incubation, image-based cell proliferation readouts followed by biochemical-based cell viability measurements were used to examine the compound effect on cancer cell growth. The percentage of control (%C) was calculated using the equation 100X(S_compound_-S_blank_)/(S_positive_-S_blank_), where S_positive_ and S_blank_ are the corresponding average of the cell fluorescence intensity for wells with DMSO containing PBMCs/medium only or plus cancer cells, respectively. The immune killing selectivity index was calculated using the equation %C_-PBMC_/%C_+PBMC_.

#### Quantitative real-time polymerase chain reaction (qPCR)

Total RNA was isolated from cell lysates using E.Z.N.A.® Total RNA Kit I (Omega, Cat# R6834-01) and digested with DNase I (Invitrogen, Cat# 18068-015). Total of 1 μg RNA was subjected to cDNA synthesis using SuperScript™ III First-Strand Synthesis System(Invitrogen, Cat# 18080051) followed the manufacture’s instruction. Reverse transcribed cDNA or isolated cytoplasmic DNA was diluted 1:5∼1:10 in nuclease free water. qPCR was performed using SYBR Green Supermix (Bio-rad, Cat# 1725272) in Mastercycler® RealPlex PCR System (Eppendorf) with primers as listed in Supplementary Table 1. All the primers were ordered from Eurofins Genomics LLC. (Louisville, KY, USA). The following thermal cycling conditions were used for IFNβ: 50 °C for 2 min; 95 °C for 10 min; 40 thermal cycles (94 °C for 10 sec, 59 °C for 30 sec, 72 °C for 45 sec and 75 °C for 29 sec) (Kotla et al., 2008). For other genes, the following thermal cycling conditions were used: 95°C for 2 min; 40 thermal cycles (95 °C for 15 sec, 60 °C for 15 sec and 72 °C for 20 sec). RNA expression was normalized to GAPDH expression. Data were conducted a comparative analysis of relative expression by 2^-ΔΔCt^ method. Primer information of qRT-PCR was shown in Supplementary Table 1.

#### Isolation of cytoplasmic dsDNA

Cytoplasmic DNA was extracted by using mitochondrial DNA isolation kit (BioVision, Cat# K280-50) according to modified manufacturer’s instructions. Briefly, 0.5 × 10^6^ cells were lysed with 1x cytosol extraction buffer, homogenized by dounce tissue grinder(50-70 times), and then the nuclei and mitochondrial fractions were removed by centrifugation according to the manufacturer’s instructions. Cytoplasmic DNA from lysate was isolated by QIAamp DNA Mini Kit(QIAGEN, Cat# 51304). Isolated cytoplasmic DNA was purified by RNaseA (Thermo Fisher Scientific, Cat# EN0531) to remove RNA contamination. The amount of mtDNA in cytosol was determined by qRT-PCR using MT-ND1 primers. The amount of nuclear DNA in cytosol was determined by qRT-PCR using three different sets of primers designed for different chromosomes as described previously (Kitajima et al., 2019). The sequences of the primers are listed in Supplementary Table 1. 40 ng DNA per well template was used for PCR analysis as described in qRT-PCR.

#### Transcriptome (RNA-seq) analysis

The birinapant-induced immune-dependent transcriptome change was analyzed by mRNA sequencing service at Novogene Corporation Inc. (Sacramento, CA,USA). Birefly, LKB1-mut cancer cells were treated by birianpant with or without PBMC co-culture for 24 hours. Then immune cells were removed by washing monolayer cancer cells three times with 1X PBS. The total RNA was isolated using E.Z.N.A.® Total RNA Kit I. RNA sequence reads were aligned to the human reference genome (GRCh38). Significantly up- or downregulated differential expression genes (DEGs) were identified using |log_2_(Fold-Change (FC))|*≥*1 and adjusted P value *≤*0.05. Pathway enrichment analysis was performed using Metascape (Zhou et al., 2019).

#### Bioinformatics analysis

The Gene Set Enrichment Analysis (GSEA) analyses was performed as described previously (Tang et al., 2021). Briefly, GSEABase package in R Studio was used to score the indicating gene sets. The Hallmark gene sets available from the Molecular Signatures database (MSigDB) (Liberzon et al., 2015) were used as the reference gene sets. The rank of genes in indicating pathways was used in accordance with the birinapant-induced immune-dependent DEGs, or DEGs identified between LKB1-WT and LKB1-mut lung adenocarcinoma patient samples from the Genomic Data Commons (GDC) Data Portal (https://portal.gdc.cancer.gov/). The normalized enrichment score (NES) was calculated to reflect the degree in which a set of genes is overrepresented at the extremes (top or bottom) among the entire ranked list. All GSEA analyses were performed strictly according to the instruction (https://www.bioconductor.org/packages/release/bioc/vignettes/GSEABase/inst/doc/GSEABase.pdf). As for statistical significance, |NES|>1 with a P value and False discovery rate (FDR) <0.05 was considered as significantly enriched.

#### Cell lysate-based affinity pulldown assay

Cell lysate-based affinity-pulldown assay was performed as we previously described (Boettcher et al., 2018; Grzeskowiak et al., 2018; Ivanov et al., 2018; Li et al., 2017; Mo et al., 2017; Mo and Fu, 2016; Mo et al., 2016; Rusnak et al., 2018; Tang et al., 2021). Briefly, cell lysate were prepared in nonidet P-40 (NP-40) lysis buffer (200 μL contains 1% NP-40 (IGEPAL CA-630, Sigma-Aldrich), 20mM Tris-HCl, 150mM NaCl, 5% glycerol and 2mM EDTA, supplemented with protease inhibitor cocktail (Sigma-Aldrich, Cat# P8340), phosphatase inhibitor cocktail 2 (Sigma-Aldrich, Cat# P5726), and phosphatase cocktail 3 (Sigma-Aldrich, Cat# P0044)). Cell lysate were subjected to affinity-based pulldown using glutathione-conjugated sepharose beads (GE, Cat# 17-0756-05) for GST-pulldown, EZview Red Anti-Flag M2 Affinity Gel (Sigma-Aldrich, Cat# F2426) for flag-immunoprecipitation, or EZview Red Anti-HA Affinity Gel (Sigma-Aldrich, Cat# E6779) for HA-immunoprecipitation. Cell lysate were incubated with beads at 4 °C for 2 hours. Beads were washed with NP-40 lysis buffer for three times. Pulldown or immunoprecipitated protein complex were eluted by boiling the beads at 95 °C for 5 minutes in 2x Laemmli buffer (Bio-rad, Cat# 1610737, supplemented with 200mM DL-Dithiothreitol (DTT)). Sample were then analyzed by SDS-polyacrylamide gel electrophoresis and immunoblotting with desired antibodies.

#### Co-immunoprecipitation (Co-IP) with endogenous proteins

The endogenous co-IP assay was performed as described previously (Mo et al., 2017; Rusnak et al., 2018; Tang et al., 2021). Briefly, cell lysate were prepared in NP-40 lysis buffer. DTT (10mM) and N-Ethylmaleimide (5mM) were added for cell lysate used for protein ubiquitination level measurement. Cell lysates with total ∼1.5mg of proteins were used for immunoprecipitation by incubating with desired protein antibody or IgG control at 4 °C for 16 hours. Then protein A/D agarose beads were added to the cell lysate and antibody mixture for incubation at 4 °C for 1 hour. Beads were then washed with NP-40 lysis buffer three times. Immunoprecipitated protein complex were eluted by boiling for 5 minutes at 95 °C in 2x Laemmli buffer without DTT. Then add DTT into supernatant at final concentration of 20mM and then boil for another 5 minutes at 95 °C. Samples were analyzed by SDS-polyacrylamide gel electrophoresis and immunoblotting with desired antibodies.

#### Time Resolved-Föster Resonance Energy Transfer (TR-FRET) assay

TR-FRET assay was performed as previously described (Li et al., 2017). The FRET buffer used throughout the assay contains 20 mM Tris-HCl, pH 7.0, 50 mM NaCl, and 0.01% NP-40. Cell lysate from HEK293T cells expressing GST-tagged and Venus-flag (VF) tagged donor and acceptor proteins were prepared in 1% NP-40 buffer. Cell lysate were serially diluted in FRET buffer and mixed with anti-GST-Terbium antibody (1:2000, Cisbio US Inc, Cat# 61GSTTLB). The plate was centrifuged at 200xg for 5 min and incubated at 4 °C for overnight. TR-FRET signals were measured using the BMG Labtech PHERAstar *FSX* reader with the HTRF optic module (excitation at 337 nm, emission A at 665 nm, emission B at 620 nm, integration start at 50 μs, integration time for 150 μs and 8 flashes per well). All FRET signals were expressed as a TR-FRET ratio: F665nm /F620nm x 10^4^.

#### Western Blot

Proteins in the SDS sample buffer were resolved by 10% SDS polyacrylamide gel electrophoresis (SDS-PAGE) and were transferred to nitrocellulose filter membranes at 100 V for 2h at 4 °C. After blocking the membranes in 5% nonfat dry milk in 1×TBST (20mM Tris-base, 150mM NaCl, and 0.05% Tween 20) for 1 hour at room temperature, membranes were blotted with the indicated antibodies at 4 °C overnight. Membranes were washed by 1×TBST for three times, 15 minutes each time. SuperSignal West Pico PLUS Chemiluminescent Substrate (Thermo, #34580) and Dura Extended Duration Substrate (Thermo, #34076) were used for developing membranes. The luminescence images were captured using ChemiDoc™ Touch Imaging System (Bio-Rad).

#### Generation of lentivirus

HEK293T cells (5×10^6^) were seeded onto a 6-well plate and transfected using PEI transfection reagent with 2 μg of pHAGE-puro-lentivirus-based expression vector together with 1.6 μg pCMV-dR8.91 and 0.66 μg of pCMV-VSVG (Ng et al., 2018). Forty-eight to 72 hours after transfection, the conditioned media containing lentivirus particles were collected and centrifuged at 4 °C, 2500xg for 15min. Then media were filtered by 0.45 μm PVDF filter (Millipore, Cat# SLHV033RS) and stored in −80 °C.

#### interferon-stimulated response element (ISRE) and Interferon Gamma-Activated Sequence (GAS) luciferase reporter assay

HEK293T cells transiently expressing GAS or ISRE-driven firefly luciferase and internal control renilla luciferase, or A549 cell stably expressing GAS or ISRE-driven firefly luciferase were used for the luciferase reporter assay. Cells were transfected with VF-tagged plasmids or treated with birinapant and/or IFN*γ* as indidated. Renilla and Firefly luciferase activities were measured by Envision Multilabel plate reader (PerkinElmer) using and Dual-Glo luciferase kit (Promega, Cat# E2920) according to the manufacturer’s instructions. The normalized luminescence was calculated as the ratio of luminescence of Firefly luciferase over the luminescence of Renilla luciferase for HEK293T cells. For stable A549 ISRE- or GAS-luciferase reporter cells, only firefly luciferase signal was measured.

#### Cytosolic double-strand DNA Staining

Cells were cultured on chambered cell culture slides (ibidi, Cat# 80826). Cells were treated with indicated treatments or vehicle control for 24 hours. Following the treatment, cells were incubated with culture media containing PicoGreen dsDNA stain (200-fold dilution) (Thermo Fisher, Cat# P7581) for 1 hour. Then cells were incubated with RPMI1640 medium (without FBS) containing 200nM MitoTracker™ Red reagent (Thermo Fisher, Cat# M7512) for 20 minutes. Then cells were fixed with 4% paraformaldehyde in PBS (Fisher scientific, Cat# 50-980-487) for 10 minutes. Cells were then washed twice with PBS and stained with Hoechst 33342 (Thermo Fisher Scientific, Cat# H3570) for 10 minutes. The image was acquired using Nikon A1R HD25 inverted confocal microscopes.

#### STING-HiBiT expression assay

STING-HiBiT engineered cells were generated by essentially following the protocol for CRISPR-mediated HiBiT tagging of endogenous proteins (Promega) (Schwinn et al., 2018). STING-HiBiT expression was monitored using Nano-Glo® HiBiT Lytic Detection System (Promega).

#### Apoptosis assay

The cell apoptosis assay was performed essentially by following manufacturer’s protocol using Incucyte® Caspase-3/7 Green Dye (SARTORIUS, Cat# 4440) (1:1000 dilution at final concentration of 5mM). Data was aquired and analyzed using IncuCyte® S3 Live-Cell Analysis System.

#### Transwell-based cell migration assay

Chemotaxis of immune cells was measured using a transwell-based cell migration assay essentially by adapting previously described methods (Justus et al., 2014). Briefly, A549 Nuclight-red cells (SARTORIUS, #4491) was plated at the bottom chamber of the transwell plate. Twenty-four hours later, cells were treated with birinapant, JAK inhibitor, or STING inhibitor as indicated followed by adding Jurkat T cells or NK92-MI cells pre-labeld with CellTracker™ Green CMFDA Dye (ThermoFisher, # C2925) at the upper chamber. Forty-eight hours later, data was acquired and analyzed using ImageXpress Micro HCS Imaging System.

#### Animal studies

The WRJ388 cell-based transplantation mouse model was used to evaluate IAP inhibitor’s anti-tumor efficacy in vivo similarly as described previously (Gilbert-Ross et al., 2017; Jin et al., 2021). Briefly, a total of 5×10^6^ exponentially growing WRJ-388 cells were transplanted into syngeneic immune-competent male KL mice through subcutaneouos injection. Male nude mice was used as immune-deficient control. Mice were randomly allocated into two groups (vehicle control and birinapant treatment, 6 and 5 mice/group for immune-competent and nude mice model, respectively). Birinapant treatment (10 mg/kg once per 3 days) began when tumor volumn reached about 300 mm^3^. Birinapant was dissolved in 12.5% (v/v) Captisol (MedChemExpress) in distilled water. At the indicated endpoint, tumor samples were harvested for further analysis.

#### Single cell mass cytometry

Immune cell profiling was performed using single cell mass cytometry as described previously (Bailur et al., 2019; Bailur et al., 2020; Bar et al., 2020; Boddupalli et al., 2016). Single cell suspensions from mouse tumor samples were stained with 16-marker panels using metal conjugated antibodies according to manufacturer-suggested protocol using Mar Maxpar® Mouse Sp/LN Phenotyping Panel Kit (Fluidigm, Cat# 201306). Cells were fixed, permeabilized, and washed according to manufacturer’s cell surface antigen staining protocol (Fluidigm). After antibody staining, cells were incubated with intercalator solution, washed, mixed with EQ Four Element Calibration Beads (catalog 201078), and acquired with mass cytometer (all reagents from Fluidigm). Gating and data analysis were performed with Cytobank (https://www.cytobank.org/). Intact viable cells were identified using cisplatin intercalator according to manufacturer-suggested concentrations (Fluidigm). viSNE analysis was performed with Cytobank.

#### IHC analysis

Five-micron-thick paraformaldehyde-fixed OCT-embedded mouse lung sections or formalin-fixed paraffin-embedded mouse tumor lung sections were used for IHC analyses as described previously (Jin et al., 2021). Slides were stained with indicated primary antibodies and horse anti-rabbit IgG (Vector) was used as the secondary antibody. DAB substrate kit was used to develop IHC signals. These samples were blinded and analyzed by a lung cancer pathologist (G.L. Sica).

#### Statistical analysis

All analyses were performed using GraphPad Prism version7.0 (GraphPad Software, La Jolla, CA). The dose-dependent PBMC-induced or small molecule induced cancer cell growth inhibition curve was established using GraphPad Prism based on the Sigmoidal dose-response (variable slope) equation. Statistical significance was assessed using student’s t test, or one way ANOVA followed by Tukey post hoc test. P-values< 0.05 were considered statistically significant.

